# Two Types of Cinnamoyl-CoA Reductase Function Divergently in Tissue Lignification, Phenylpropanoids Flux Control, and Inter-pathway Cross-talk with Glucosinolates as Revealed in *Brassica napus*

**DOI:** 10.1101/2021.03.01.433400

**Authors:** Nengwen Yin, Bo Li, Xue Liu, Ying Liang, Jianping Lian, Yufei Xue, Cunmin Qu, Kun Lu, Lijuan Wei, Rui Wang, Jiana Li, Yourong Chai

## Abstract

Cinnamoyl-CoA reductase (CCR) is the entry point of lignin pathway and a crucial locus in dissection and manipulation of associated traits, but its functional dissection in Brassicaceae plants is largely lagged behind though *Arabidopsis thaliana CCR1* has been characterized to certain extent. Here, 16 *CCR* genes are identified from *Brassica napus* and its parental species *B. rapa* and *B. oleracea*. Brassicaceae *CCR* genes are divided into *CCR1* subfamily and *CCR2* subfamily with divergent organ-specificity, yellow-seed trait participation and stresses responsiveness. *CCR1* is preferential in G- and H-lignins biosynthesis and vascular development, while *CCR2* has a deviation to S-lignin biosynthesis and interfascicular fiber development. *CCR1* has stronger effects on lignification-related development, lodging resistance, phenylpropanoid flux control and seed coat pigmentation, whereas *CCR2* controls sinapates levels. *CCR1* upregulation could delay bolting and flowering time, while *CCR2* upregulation weakens vascular system in leaf due to suppressed G lignin accumulation. Besides, *CCR1* and *CCR2* are deeply but almost oppositely linked with glucosinolates metabolism through inter-pathway crosstalk. Strangely, upregulation of both *CCR1* and *CCR2* could not enhance resistance to UV-B and *S. sclerotiorum* though *CCR2* is sharply induced by them. These results provide systemic dissection on *Brassica CCR*s and *CCR1*-*CCR2* divergence in Brassicaceae.

**Highlight:** Brassicaceae contains two types of Cinnamoyl-CoA reductase. As revealed in *Brassica napus*, they are divergently involved in lignin monomer biosynthesis, tissue lignification, phenylpropanoid flux control, and inter-pathway crosstalk with glucosinolates.

## Introduction

Lignins constitute one important group of phenylpropanoids, are deposited in plant secondary cell walls, and are the second most abundant biopolymers on the planet (Huang *et al*., 2010; Vanholme *et al*., 2010). They perform many functions, providing structural support, giving rigidity and strength to stems to stand upright, and enabling xylems to withstand the negative pressure generated during water transport (Escamilla-Treviño *et al*., 2010; Labeeuw *et al*., 2015). In addition, lignins have been suggested to be induced upon various biotic and abiotic stresses, such as pathogen infection, insect feeding, drought, heat and wounding (Caño-Delgado *et al*., 2003; Weng and Chapple, 2010; Chantreau *et al*., 2014).

To date, engineering of lignins mainly pursues reduced lignin content or altered lignin composition to meet the demands of agro-industrial processes, such as chemical pulping, forage digestibility, and the bioethanol production from lignocellulosic biomass (Baucher *et al*., 2003; Li *et al*., 2008; Weng *et al*., 2010; Ko *et al*., 2015; De Meester *et al*., 2020). However, when altering the expression of lignin pathway genes, the metabolic flux of its neighbor pathways would be changed correspondingly (Hoffmann *et al*., 2004; Li *et al*., 2010; Thévenin *et al*., 2011). Furthermore, dramatic modification of lignin content or lignin composition may provoke deleterious effects on plant growth, such as dwarfism and collapsed xylem vessels, with concomitant loss of biomass and yield (Piquemal *et al*., 1998; Ruel *et al*., 2009; De Meester *et al*., 2020).

Cinnamoyl-CoA reductase (CCR) is the entry point for the lignin-specific branch of the phenylpropanoid pathway and catalyzes the monolignol biosynthesis (Lacombe *et al*., 1997; Kawasaki *et al*., 2006). *Arabidopsis thaliana* possesses 11 annotated *CCR* homologs (Costa and Dolan, 2003), but only *AtCCR1* and *AtCCR2* encode true CCR enzyme (Lauvergeat *et al*., 2001). *AtCCR1* is preferentially expressed in tissues undergoing lignification, while *AtCCR2* is poorly expressed during development but is strongly and transiently induced by *Xanthomonas campestris*, suggesting that *AtCCR1* might be involved in constitutive lignification whereas *AtCCR2* might be involved in resistance (Lauvergeat *et al*., 2001). However, to date there is no functional verification of this assumption through over-expression transgenic study in *A. thaliana*, and the function of *CCR2* in Brassicaceae is not dissected. In monocot grasses, *CCR1* expression can be detected in various organs with a relatively high transcription level in stem (Tu *et al*., 2010; Giordano *et al*., 2014), thus is thought to be involved in constitutive lignification. In poplar and switchgrass, *CCR2* is expressed at very low levels in most organs, but can be significantly induced by biotic and abiotic stresses (Escamilla-Treviño *et al*., 2010). Manipulation of *CCR* (mainly through downregulation) typically results in a significant variation of lignin content and composition (Goujon *et al*., 2003; Zhou *et al*., 2010; Wagner *et al*., 2013). Moreover, plants with dramatically downregulated *CCR* genes usually showed a stunted growth and delayed development (Tamasloukht *et al*., 2011; Thévenin *et al*., 2011; De Meester *et al*., 2020), and the alternation of carbon flux between lignin and other metabolic pathways was also accompanied (van der Rest *et al*., 2006; Dauwe *et al*., 2007; Wagner *et al*., 2013).

Lodging and diseases are fatal problems in field production of most crops. Reducing plant height has been proven to be a useful strategy for improving lodging resistance, but dwarfism will reduce canopy photosynthetic capacity and yield (Zhang *et al*., 2001; Acreche and Slafer, 2011; Peng *et al*., 2014). Increasing stem strength would be a very promising strategy for breeding crops with high lodging resistance (Ma, 2009). Moreover, lignin is a physical barrier which can restrict pathogens to the infection site and confer disease resistance in plants (Lee *et al*., 2019). There are few reports at the molecular level to address how the regulation of lignin biosynthesis affects crop lodging and pathogen resistance via gene manipulation. In wheat, accumulation of lignin is closely related to lodging resistance, and wheat culms with higher lignin content could have a better lodging resistance (Peng *et al*., 2014). Regarding defense, the *CCR*-like gene *Snl6* is required for NH1-mediated resistance to bacterial pathogen *Xanthomonas oryzae* pv. *Oryzae* in rice (Bart *et al*., 2010). However, there is few reports on overexpressing *CCR* in enhancing lodging or disease resistance with molecular breeding values in crops.

*Brassica*, a relative genus of model plant *A. thaliana*, is of great importance since it contains various important oilseed, vegetable and ornamental crops. Unfortunately, *Brassica* crops especially rapeseed (*B. napus*) frequently suffer from some detrimental stresses such as lodging (Liu *et al*., 2010; Peng *et al*., 2014) and stem rot disease caused by *Sclerotinia sclerotiorum* with serious yield penalty and quality deterioration (del Río *et al*., 2007; Ding *et al*., 2015). In rapeseed, lodging could lead to 20%–46% yield loss and about four percent points of oil content reduction, and limits the efficiency of mechanical harvest (Pan *et al*., 2012; Kendall *et al*., 2017; Berry, 2018). There is little knowledge at present concerning the functional genes involved in lignin biosynthesis in rapeseed or even in *Brassica* species. Comprehensive characterization of the lignin biosynthesis in rapeseed will enable us to manipulate the lignin content or composition through genetic engineering, and to strengthen the capacity for lodging and pathogen resistance.

In this study, the *CCR1* and *CCR2* subfamily members from *B. napus* and its parental species *B. rapa* and *B. oleracea* were isolated, their expression patterns in different organs and under various stresses were investigated, and overexpression of *BnCCR1* and *BnCCR2* was performed for deciphering their biological functions and biotechnological potentials. Our results demonstrated that *BnCCR1* was mainly involved in the biosynthesis of H- and G-lignins, while *BnCCR2* showed a preference in the biosynthesis of S-lignin and in plant defense. A dramatic shift of carbon flux in phenylpropanoid pathway and a strong crosstalk effect on glucosinolate pathway was demonstrated in both *BnCCR1-* and *BnCCR2-*transgenic plants, especially for *BnCCR1* manipulation. Besides, *BnCCR1* and *BnCCR2* showed distinctly different association with flux regulation and development control, which accounted for distinct phenotypic modification in vascular system, lodging resistance, seed color, flowering time and glucosinolate profiles in corresponding overexpressors. Surprisingly and unexpectedly, both *BnCCR1-* and *BnCCR2-*overexpressors did not show increased resistance to *S. sclerotiorum* and UV-B light, implying complicated association of *CCR* and lignin pathway with disease resistance.

## Materials and methods

### Plant Materials

*B. napus*: black-seed varieties 5B and ZS10, yellow-seed variety 09L587. *B. rapa*: black-seed variety 09L597 and yellow-seed variety 09L600. *B. oleracea*: black seed variety 09L598 and yellow-seed variety 09L599. T1 transgenic lines together with non-transgenic wild-type (WT) ZS10 plants were grown in artificial growth room (25°C, 16-h photoperiod/20°C, 8-h dark period). Later generations of transgenic lines and all other materials were planted in field cages, with common plantation conditions. Samples were immediately frozen in liquid nitrogen and stored at −80°C for gene expression analysis, biochemical and histochemical assay, gas chromatography–mass spectrometry (GC-MS) detection and UPLC-HESI-MS/MS analysis. Mature seeds, stems and roots (about 65 DAP) were harvested for agronomic traits investigation and biochemical analysis.

### Southern hybridization analysis

Total genomic DNA was extracted from leaves of the 3 *Brassica* species using a standard cetyltrimethylammonium bromide protocol (Porebski *et al*., 1997). It was digested with restriction enzymes *Dra*I, *Eco*RI, *Eco*RV, *Hin*dIII and *Xba*I (70 µg for each enzyme) respectively, and separated on a 0.8% (w/v) agarose gel. Following electrophoresis, DNA was transferred to a positively charged nylon membranes (Roche, Switzerland) using established protocols. The *Brassica CCR1*- and *CCR2*-specific probes were amplified by PCR with primer pairs FCCR1C1+RCCR1 and FBCCR2I+RBCCR2I respectively, and labelled using digoxigenin (DIG) Probe Synthesis Kit (Roche, Switzerland), with an annealing temperature of 61°C and extension time of 1 min. Sequences of all primers are provided in Supplementary Table S8. Hybridizations with the probes were performed at 43°C overnight, with chemiluminescent detection using DIG Luminescent Detection Kit (Roche, Switzerland).

### Detection of Transcription Levels

Total RNA of each sample was extracted using EASYspin Kit (Biomed, China) and RNAprep pure plant kit (TIANGEN, China), and treated with DNase I to eliminate contaminated gDNA. Equal quantities of RNA (1µg) were adopted for the synthesis of total cDNA using the PrimeScript^TM^ RT reagent kit with gDNA Eraser (TaKaRa Dalian, China). The transcript levels of *CCR1*, *CCR2* and other target genes were detected using both quantitative real-time PCR (qRT-PCR) and semi-quantitative RT-PCR (sRT-PCR) as described previously (Zhao *et al*., 2007b). The *25SrRNA* primer pair was used as an internal control in qRT-PCR. A 25-fold dilution series of original reverse-transcription products was used for qRT-PCR using SsoAdvaned^TM^ Universal SYBR Green Supermix (BioRad, USA) on CFX96^TM^ Real-Time System (BioRad, USA). Conditions for qRT-PCR were as follows: 95°C for 2 min; 40 cycles of amplification with 95°C for 30 s and 62°C for 30 s. A melting curve was obtained after amplification by heating products from 60 to 95°C. Transcript levels were determined based on changes in Cq values relative to the internal control. Results were analyzed using the CFX Manager^TM^ 3.0 software (Bio-Rad, USA). Conditions for sRT-PCR on Veriti Thermal Cycler (ABI, USA) were as follows: 94°C for 2 min; 31 cycles of amplification with 94°C for 0.5 min, 60-64°C for 0.5 min and 72°C for 1 min; followed by 72°C for 10 min. The *26SrRNA* primer pair was used as an external control in sRT-PCR. Sequences of all primers for qRT-PCR and sRT-PCR are listed in Supplementary Table S8.

### Rapeseed Genetic Transformation and Screening

A modified *Agrobacterium*-mediated transformation protocol according to (Cardoza and Stewart, 2003) was used to transform “double low” (low erucic acid and low glucosinolates) rapeseed commercial cultivar Zhongshuang 10 (ZS10) with overexpression vectors pCD-*BnCCR1-2ox* and pCD-*BnCCR2-4ox*, using hypocotyl segments as explants (Fu *et al*., 2017). The regenerated plants were identified first by leaf β-glucuronidase (GUS) staining and leaf Basta-resistance test (200 ppm), then by Taq-PCR detection of target genes (as those for engineering strains verification) and the selectable and screenable marker gene *BAR* (primer pair FBar+RBar, annealed at 58°C, extension for 30 s). The T1 transgenic plants were grown in artificial growth room and were selfed. Representative T2 and T3 transgenic lines and WT were grown in field cages, and positive plants of above identifications were subjected to traits investigation and further studies.

### Agronomic Traits Investigation

The area and length of the 4th or the 5th leaves which was fully expanded were surveyed during vegetative stage, and the length and width of the petioles of these leaves were measured at the same time. The primary branch numbers, middle stem diameter and stem strength at reproductive stage, and the plant traits at harvest stage, were recorded and measured. The weight of 1000 seeds and yield per plant were measured after the harvested seeds were dry. Stem strength determination: Freshly collected middle stem segments were placed horizontally, and the force exerted to break the stem was recorded with a universal force testing device (model DC-KZ300, Sichuan, China) to determine the stems rigidity, which was normalized with the stem’s length and diameter.

### Quantification of Insoluble Condensed Tannins of Seed Coat

The modified method for quantification of insoluble condensed tannins was according to (Auger *et al*., 2010) and (Naczk *et al*., 2000). The oven-dried seed coat was milled to fine powder using a microball mill, and extracted with hexane for 12 h using a Soxhlet apparatus and then dried at room temperature. 10 mg sample of seed coat powder was extracted with the extraction solution of three milliliters of butanol-HCl (95:5; v/v), 300 μL methanol and 100 μL of 2% ferric ammonium sulfate (w/v) in 2 N HCl. The tubes were heated for 3 h at 95°C in a water bath, centrifuged after cooling, and extracted again with the same extraction solution for 1 h. The absorbance of the pooled supernatant was measured at 550 nm against a reagent-only blank using UV-VIS spectrophotometer (UV-5100B, Shanghai, China). A calibration curve was prepared using procyanidin with amounts ranging from 0 to 200 μg/mL.

### Histochemical Staining, Autofluorescence and Anatomical Studies

Cross sections were obtained by using a frozen section machine (Leica CM1850, Germany). Fresh petioles at vegetative stage, fresh stems and roots of the plants at reproductive stage, fresh mature silique coat and dry mature seeds coat were cut into slices of 60, 60, 60, 60 and 5 μm thick, respectively. Phloroglucinol-HCl staining and Mäule staining of those plant organ sections were performed as previously described (Chapple *et al*., 1992; Chen *et al*., 2002). Stained sections were observed on a stereoscopic microscope (Nikon C-BD230, Japan; OLYMPUS SZX2-FOA, Japan) and a fluorescence microscope (Nikon ECLIPSE E600W, Japan).

### Determination of Lignin Content and Composition

Lignin content was quantified using a modified acetyl bromide soluble lignin method (Fukushima and Hatfield, 2001; Chang *et al*., 2008), and lignin composition was determined with a modified thioacidolysis method according to previous reports (Lapierre *et al*., 1995; Yosef and Ben-Ghedalia, 1999; Robinson and Mansfield, 2009).

#### Cell Wall Isolation

Harvested stems were dried in a 70°C forced air oven for 72 h, and then were ground with a Wiley mill to pass a 40-mesh screen or with a microball mill to pass an 80-mesh screen using to meet the requirements of the subsequent extraction method. 60-70 mg samples were added into a 2 mL Sarstedt screw cap tube, adding with 1.5 mL of 80% aqueous ethanol to the dispensed ground material, and vortex thoroughly (10 min). This step was repeated for a total of three cycles, followed by centrifugation at 10,000 rpm for 10 min, and the supernatant was decanted. Add 1.5 mL of chloroform/methanol (2:1 v/v) solution to the residue, and shake the tube thoroughly to resuspend the pellet. Centrifuge at 10,000 rpm for 10 min, and decant the supernatant. Resuspend the pellet in 500 μL of acetone, and evaporate the solvent with a stream of air at 35°C until dry (If needed, dried samples can be stored at room-temperature until further processing). To initiate the removal of starch from the sample, resuspend the pellet in 1.5 mL of a 0.1 M sodium acetate buffer pH 5.0, cap the Sarstedt tubes, and heat for 30 min at 80°C in a heating block. Cool the suspension on ice, and add the following agents to the pellet: 35 μL of 0.01% sodium azide (NaN_3_), 35 μL amylase (from Bacillus species, 50 μg/mL in H_2_O, Sigma); 3.56 μL pullulanase (from Bacillus acidopullulyticus, 17.8 units, Sigma). Cap the tube, and vortex thoroughly. The suspension is incubated over night at 37°C in the shaker (Orienting the tubes horizontally aides improved mixing). After enzyme digestion, the suspension was heated at 100°C for 10 min in a heating block to terminate digestion. Centrifuge at 10,000 rpm for 10 min, and discard the supernatant containing solubilized starch. The remaining pellet was washed three times by adding 1.5 mL water, vortexing, centrifuging, and decanting of the washing water. For removing the water, 1.5 mL anhydrous ethanol was added to resuspend the pellet, followed by vortexing, centrifuging, decanting of the supernatant, and resuspending the pellet with 500 μL of acetone. Finally, evaporate the solvent with a stream of air at 35°C until dry. It may be necessary to break the material in the tube with a spatula for better drying. If needed, the dried samples can be stored at room-temperature until further processing.

#### Spectrophotometer Test of Lignin Content

2 mg of prepared cell wall material was interacted with 200 μL of freshly made acetyl bromide solution (25% v/v acetyl bromide in glacial acetic acid) in a culture tube under 50°C for 3 h with shaking at 30 min intervals during first 2 h and with vortex every 15 min during the last 1 h, and the reaction was stopped on ice for 5 min. Add 800 μL of 2 M sodium hydroxide and 140 μL of freshly prepared 0.5 M hydroxylamine hydrochloride, and vortex the tubes. Transfer the reaction solution to a 15 mL graduated test tube with stopper, and the tubes were rinsed with glacial acetic acid to complete the transfer. Fill up the tubes exactly to the 15.0 mL mark with glacial acetic acid, cap, and invert several times to mix. The absorbance of the solutions was read at 280 nm on a Varian Cary 50 spectrophotometer. A blank was included to correct for background absorbance by the reagents. Determine the percentage of acetyl bromide soluble lignin (%ABSL) using the coefficient (23.35) with the following formula: 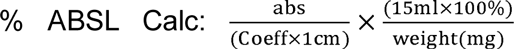, 1 cm represents the pathlength, multiplication of %ABSL with 10 results in the ug/mg cell wall unit.

#### GC-MS Test of Lignin Composition

Transfer approximately 5 mg of cell wall material into a screw-capped glass tube for thioacidolysis. Add 1 mL freshly made thioacidolysis reagent consisting of 2.5% boron trifluoride diethyl etherate (BF3) and 10% ethanethiol (EtSH) in dioxane solution, purge vial headspace with nitrogen gas (N_2_), and cap immediately. After 4 h reaction at 100°C with gentle mixing every hour, the reaction was stopped by cooling on ice for 5 min. The pH of the reaction was adjusted to 3-4 by adding 300 μL of 0.4 M sodium bicarbonate. Add 2 mL water, 200 μL tetracosane (0.5 mg/ml ethyl acetate) and 0.8 mL of ethyl acetate, and vortex. Transfer 300 μL of the ethyl acetate layer into a 2 mL Sarstedt tube (make sure no water is transferred), and evaporate the solvent with N_2_. To remove excess water, 200 μL acetone was added and evaporated twice. The finally obtained oily residue was redissolved in 500 μL of ethyl acetate, and 100 μL of resuspended sample was added with 20 μL of pyridine, and 100 μL of N, O-bis(trimethylsilyl) acetamide for the TMS derivatization, and incubated for 2 h at 25°C. The GC-MS test was performed to identify and quantify the lignin monomer trimethylsilylated derivatives, as 1 μL injecion volume sample was separated with a Restek Rxi-5ms column (30 m X 0.25 mm X 0.25 μm film thickness) under split mode, and the injector and detector temperatures were set to 250°C. Helium was the carrier gas. The following temperature gradient was used with a 30 min solvent delay and a 1.1 mL/min flow rate: Initial hold at 130°C for 3 min, a 3°C/min ramp to a 250°C whish was hold for 5 min, and then equilibration to the initial temperature of 130°C. Quantitation of the main lignin-derived monomers was performed after an appropriate calibration relative to the Tetracosane internal standard, and the characteristic mass spectrum ions of 299 m/z, 269 m/z and 239 m/z were representative for S, G and H trimethylsilylated derivatives respectively.

### Phenolic Profiling by UPLC-HESI-MS/MS

#### Extraction of Soluble Metabolites

Samples of stems, leaves, petioles and seeds (30 DAP) were ground under liquid nitrogen in a mortar and pestle, and freeze-dried by using a vacuum freeze drier (SCANVAC, Coolsafe 110 – 4, Denmark). 30 mg ground lyophilized stems, leaves and petiole material were extracted twice by sonication with 1.0 mL of 50% methanol plus 1.5% acetic acid for 1 h at 4°C, clarified at 15,000×g for 10 min. The supernatants were combined and concentrated by using a vacuum concentration (SCANVAC, scan speed 32, Denmark), and redissolved in 0.5 mL of 50% methanol, filtered through a 0.22 µm nylon syringe filter. 50 mg ground lyophilized seed material was extracted as previously described (Auger *et al*., 2010) with a little modification. One ml of a methanol/acetone/water/TFA mixture (40:32:28:0.05, v/v/v/v) was added to the seed samples, and sonicated for 1 h at 4°C. After centrifugation (15,000 ×g, 5 min), the pellet was extracted further with 1 mL methanol/acetone/water/TFA mixture overnight at 4°C under agitation (200 rpm), while the supernatant was stored at −80°C. Supernatants were pooled and clarified at 15,000 ×g for 10 min, then concentrated. To further remove the water, 200 μL methanol was added to the extracts two times during evaporation. The dried extracts were redissolved in 1 mL of 1% acetic acid in methanol and filtered through a 0.22 µm nylon syringe filter, then stored at −80°C before analysis.

#### UPLC-HESI-MS/MS Analysis

UPLC was performed on Dionex Ultimate-3000 UHPLC System (Thermo Fisher Scientific, Tacoma, Washington, USA), as 5 μL samples were separated on a Waters ACQUITY UPLC BEH C18 column (1.7 µm, 2.1 mm ×150 mm). The flow rate was 0.2 mL/min, and the oven temperature was 30°C. Eluent A was 0.1% formic acid in water, and eluent B was 0.1% formic acid in acetonitrile. The following gradient was applied for stems, leaves and petiole extracts elution: 5% B for 5 min, 5% B to 95% B for 20 min, 95% B for 5 min, followed by column wash and re-equilibration. For seed metabolites elution, gradient conditions were as follows: 5% to 9% B for 5 min, 9% B to 16% B for 10 min, 16% B to 50% B for 25 min, 50% B to 95% B for 15 min, 95% B for 5 min, followed by column wash and re-equilibration. Mass analyses were carried out with the mass spectrometer Thermo Scientific™ Q Exactive™ (Thermo Fisher Scientific, Tacoma, Washington, USA) equipped with a HESI source used in the negative ion mode. Source parameters were as following: spray voltage of 3.0 kV, sheath gas flow rate at 35 arbitrary units, auxiliary gas flow rate at 10, capillary temperature at 350°C, and aux gas heater temperature at 300°C. Nitrogen gas was used as sheath gas and auxiliary gas. Full MS/dd-MS2 were acquired from m/z 100 to m/z 1500. Thermo Xcalibur software version 3.0.63 (Thermo Fisher Scientific) was used for data collection and processing. Contents of the metabolites were expressed relative to the calibration curves of available standards. Standard compounds, namely *p*-coumaric acid, caffeic acid, ferulic acid, epicatechin, quercetin, kaempferol and isorhamnetin (from PureChem-Standard Co., Ltd, Chengdu, China) as well as sinigrin, sinapic acid, coniferyl aldehyde and abscisic acid (from Sigma-Aldrich, USA) were analyzed under the same conditions described above.

### Statistical Issues

Except for non-necessary cases and specially indicated situation, all experiments in this study were carried out with 3 replications, and all experimental results were statistically analyzed. Statistical significance was calculated by two-tailed Student’s t test (*P <0.05, **P < 0.01), and error bars indicate SD. One-way ANOVA followed by Duncan’s multiple comparisons test was used to determine the differences. Values of P < 0.05 were considered to be statistically significant.

### Accession Numbers

Sequence data of the genes and proteins involved in this article can be found in the NCBI (http://www.ncbi.nlm.nih.gov/) databases under the accession numbers indicated in Supplementary Table S1 and Fig. 1.

**Fig. 1.**
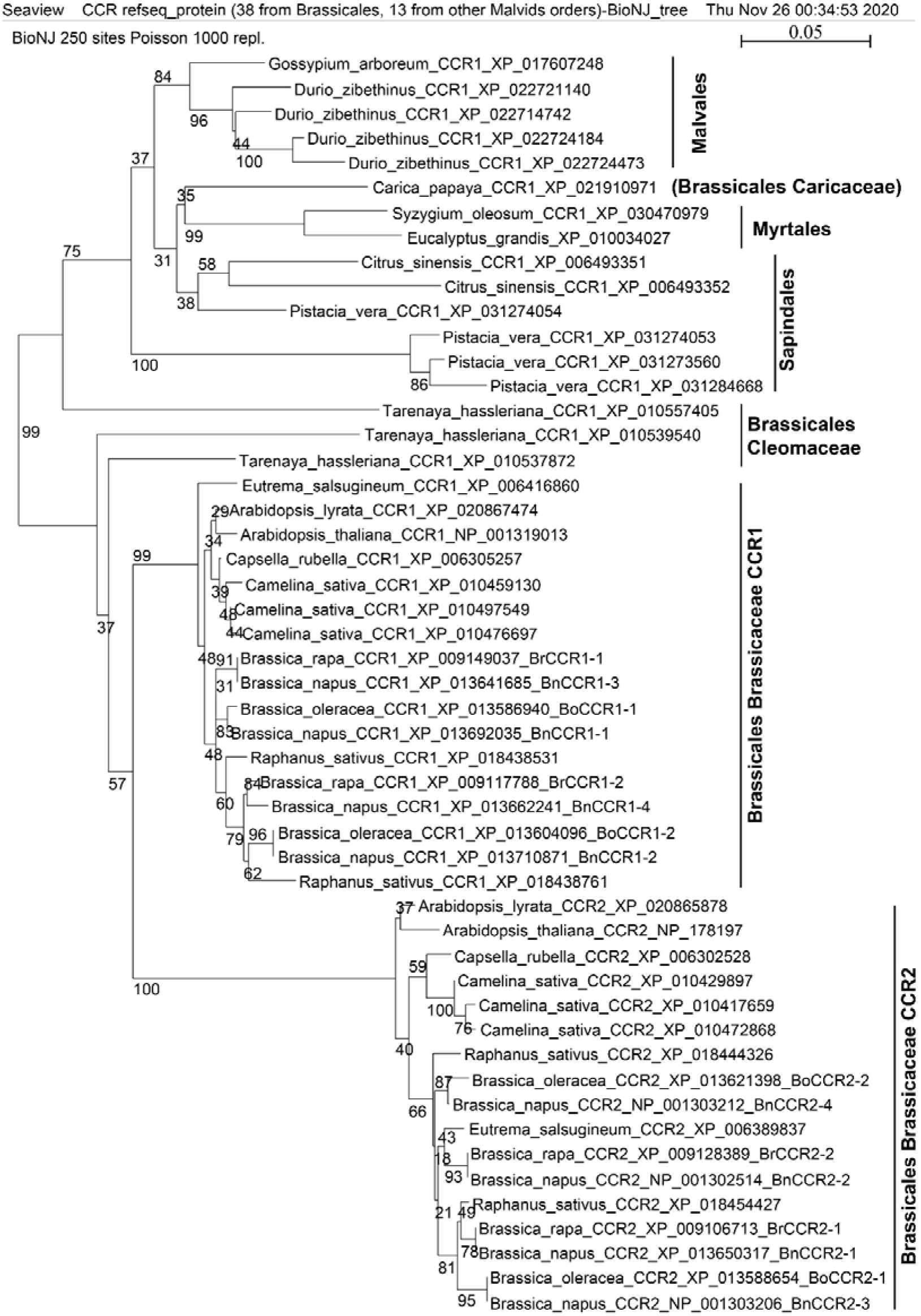
Phylogenetic analysis of CCR refseq_proteins from representative whole-genome-sequenced Malvids species including *B. napus, B. rapa* and *B. oleracea*. Latin name, refseq_protein name and accession number are provided for each sequence. Names of cloned genes in this study are given after the accession numbers. Vertical bars on the right indicate corresponding order or order-family names.

### Other Methods

Details of the stress treatments, bioinformatic methods for this study, construction of overexpression vectors, measurement of leaf spad readings, and near-infrared reflectance spectroscopy (NIRS) measurements are provided in supplementary methods.

## Results

### Isolation and characterization of *CCR* genes from *Brassica* species

*CCR* genes were isolated from *B. napus* and its parental species *B. rapa* and *B. oleracea* using RACE strategy. Full-length cDNAs and corresponding gDNA sequences of three, four and six *CCR1*-subfamily gene sequences were isolated from *B. rapa*, *B. oleracea* and *B. napus*, while three, three and four *CCR2*-subfamily gene sequences were isolated from *B. rapa*, *B. oleracea*, respectively. Besides, one *CCR2* pseudogene from each parental species and two *CCR2* pseudogenes from *B. napus* were also isolated (Supplementary Table S1). To determine the copy number of the *BnCCR1*, *BoCCR1*, *BrCCR1*, *BnCCR2*, *BoCCR2* and *BrCCR2* genes, Southern hybridization analysis was performed (Supplementary Fig. S1), and the numbers of the clear bands were well correlated with the cloned gene sequence numbers. Supplementary Table S1 shows all identity parameters of *CCR1*-subfamily and *CCR2*-subfamily genes/pseudogenes from *B. napus* and its parental species *B. oleracea* and *B. rapa* in terms of our cloning work and three genome datasets (NCBI GenBank, Genoscope and BRAD). The basic features of the cDNA and gDNA sequences are displayed in Supplementary Table S2, and the deduced protein features are depicted in Supplementary Table S3.

Multiple alignments showed that the conserved NADP binding domain and the CCR-featured traditional motif (KNWYCYGK, which was believed as the catalytic site) and novel motif H_202_K_205_R_253_ (CCR-SBM or CCR substrate binding motif) are conserved in all *Brassica* CCRs (Lacombe *et al*., 1997; Chao *et al*., 2019). Therefore, it is speculated that these *Brassica* CCRs have catalytic activities. A phylogenetic tree was constructed to reveal the relationships of *Brassica* CCR1 and CCR2 proteins with CCRs from all whole-genome-sequenced species of other Brassicales species and other malvids orders (Supplementary Fig. S2). Firstly, it is clear that an early intrafamily duplication event generated the CCR1 and CCR2 groups within Brassicaceae, i.e. AtCCR1 and AtCCR2 have respective CCR1 and CCR2 orthologs only within Brassicaceae family. Secondly, in Brassicaceae the CCR1 group is much more conserved than the CCR2 group, which could be reflected by the great difference in branch lengths. Thirdly, outside the Brassicaceae family, other Brassicales families (e.g. Cleomaceae and Caricaceae) and other malvids orders (e.g. Malvales, Myrtales and Sapindales) have similar trends in CCR evolution (gene duplication and unequal divergence of paralogs) as revealed in Brassicaceae. However, paralog numbers (numbers of duplication events) vary distinctly among relative families and orders, and the single-copy CCR from Brassicales species *Carica papaya* is nearer to non-Brassicales CCRs than to other Brassicales CCRs.

### Divergent involvement in organ-specificity and seed coat color among *Brassica CCR* family members

In all the three *Brassica* species the highest *CCR1* subfamily expression was detected in silique pericarp, while the highest *CCR2* subfamily expression varied among species (bud in *B. napus*, silique pericarp in *B. rapa*, and root in *B. oleracea*), implying faster divergence of organ-specificity in *CCR2* than in *CCR1* among species. Within each species, divergence of organ-specificity and expression intensity among *CCR* paralogs were distinct, and generally stronger divergence of organ-specificity among *CCR2* paralogs could be observed than among *CCR1* paralogs. For *CCR1* subfamily, *BnCCR1-2*, *BrCCR1-2* and *BoCCR1-1* were the dominant paralog within respective species. For *CCR2* subfamily, *BnCCR2-4*, *BrCCR2-1A* and *BoCCR2-2* were the dominant paralog within respective species (Supplementary Figs S3-S6). In all the three species, *CCR1* subfamily overall expression was distinctly lower in developing seeds especially in late-stage seeds of yellow-seed stocks than in black-seed stocks, whereas *CCR2* subfamily showed an opposite trend (Supplementary Figs S3-S6).

### Differential responsiveness to various stresses between *BnCCR1* and *BnCCR2* subfamilies

For *BnCCR1*, its overall expression showed limited upregulations by NaCl, drought & high temperature and UV-B treatment, while kept almost constant when treated with other stresses (Supplementary Fig. S7). Under the same stresses, both the overall and the member-specific expressions of *BnCCR2* subfamily were dramatically upregulated. Its overall expression in leaves was upregulated by 400.3 folds at 48 h after *S. sclerotiorum* inoculation, by 49.59 folds at 80 min after UV-B treatment, by 10.75 folds at 48 h after *P. rapae* inoculation, and with slow and slight increase after high temperature & drought treatment (Supplementary Figs S7 and S8). All *BnCCR2* subfamily members could be triggered by these stresses, and *BnCCR2-4* was dominant among paralogs (Supplementary Fig. S8). Although both *BnCCR1* and *BnCCR2* subfamilies can be distinctly regulated by multiple stresses, *BnCCR1* only mildly responds to abiotic stresses, while *BnCCR2* responds sharply and intensively to both biotic and abiotic stresses, thus *BnCCR2* is speculated to play more important roles than *BnCCR1* in coping with various stresses in *B. napus*.

### *BnCCR* over-expression dramatically influenced plant morphology, but did not increase resistance to UV-B and *S. sclerotiorum*

To further uncover the function of *Brassica CCR* genes, overexpression transgenic plants were generated. *BnCCR1-2* and *BnCCR2-4* were selected for transgenic study as they are dominant members within respective subfamilies. The transgenic lines were coded as *BnCCR1-2ox* or ox1 lines for *BnCCR1-2* overexpression, and *BnCCR2-4ox* or ox2 lines for *BnCCR2-4* overexpression, in the following analysis. The transgenic lines were screened along with non-transgenic controls (WT) by GUS staining, Basta resistance and PCR detection (Supplementary Fig. S9). Six *BnCCR1-2ox* and seven *BnCCR2-4ox* triple-positive lines were obtained, with different over-expression levels revealed by qRT-PCR detection (Supplementary Fig. S10).

All *BnCCRox* plants showed a stronger morphological development than WT throughout the whole life (Fig. 2). Both *BnCCR1-2ox* and *BnCCR2-4ox* had larger and longer leaves, higher leaf chlorophyll content, larger stem diameter, wider silique, higher breaking-resistant stem strength, better lodging resistance and more siliques per plant, with distinctly stronger effects in *BnCCR1-2ox* lines than in *BnCCR2-4ox* lines (Fig. 2; Supplementary Figs S11 and S12). Phytohormone detection showed that abscisic acid (ABA) increased significantly in the leaves of *BnCCRox* lines compared with WT (Supplementary Fig. S13). The leaves of *BnCCR1-2ox* had more obvious wrinkles and less leaf margin serrates than WT (Fig. 2c; Supplementary Fig. S12E). *BnCCR2-4ox* plants had a looser morphology with larger leaf angles (Fig. 2B and J), and their leaves were more easily to show bending and rollover phenomenon than WT when under strong sunlight and high temperature conditions (Supplementary Fig. S12D). On the other hand, upper stems of *BnCCR1-2ox* plants at late bolting stage were more easily to appear bending phenomenon, but this phenomenon would disappear after flowering stage (Fig. 2F-I; Supplementary Fig. S12A). Moreover, *BnCCR1-2ox* lines flowered 7-10 days later than WT on average, and this phenomenon varied among years and environments. However, *BnCCR2-4ox* plants had no significant difference in flowering time when compared with WT. Interesting phenomenon was observed on petiole-vein system, which was distinctly larger in transgenic plants than WT, but phloroglucinol-HCl staining showed that petiole-vein lignification was strengthened only in *BnCCR1-2ox* plants while was weakened in *BnCCR2-4ox* plants (Fig. 2D; Supplementary Fig. S12F, H and J). Seed yield per plant of *BnCCR1-2ox* had little difference compared with WT, but *BnCCR2-4ox* had slightly decreased seed yield (Supplementary Fig. S11L). Besides, both *BnCCR1-2ox* and *BnCCR2-4ox* plants exhibited no better resistance to *S. sclerotiorum* inoculation and UV-B treatment when compared with WT, even some transgenic lines had a little reduction in disease resistance in leaf identification (Supplementary Fig. S14A, B, E and H).

**Fig. 2.**
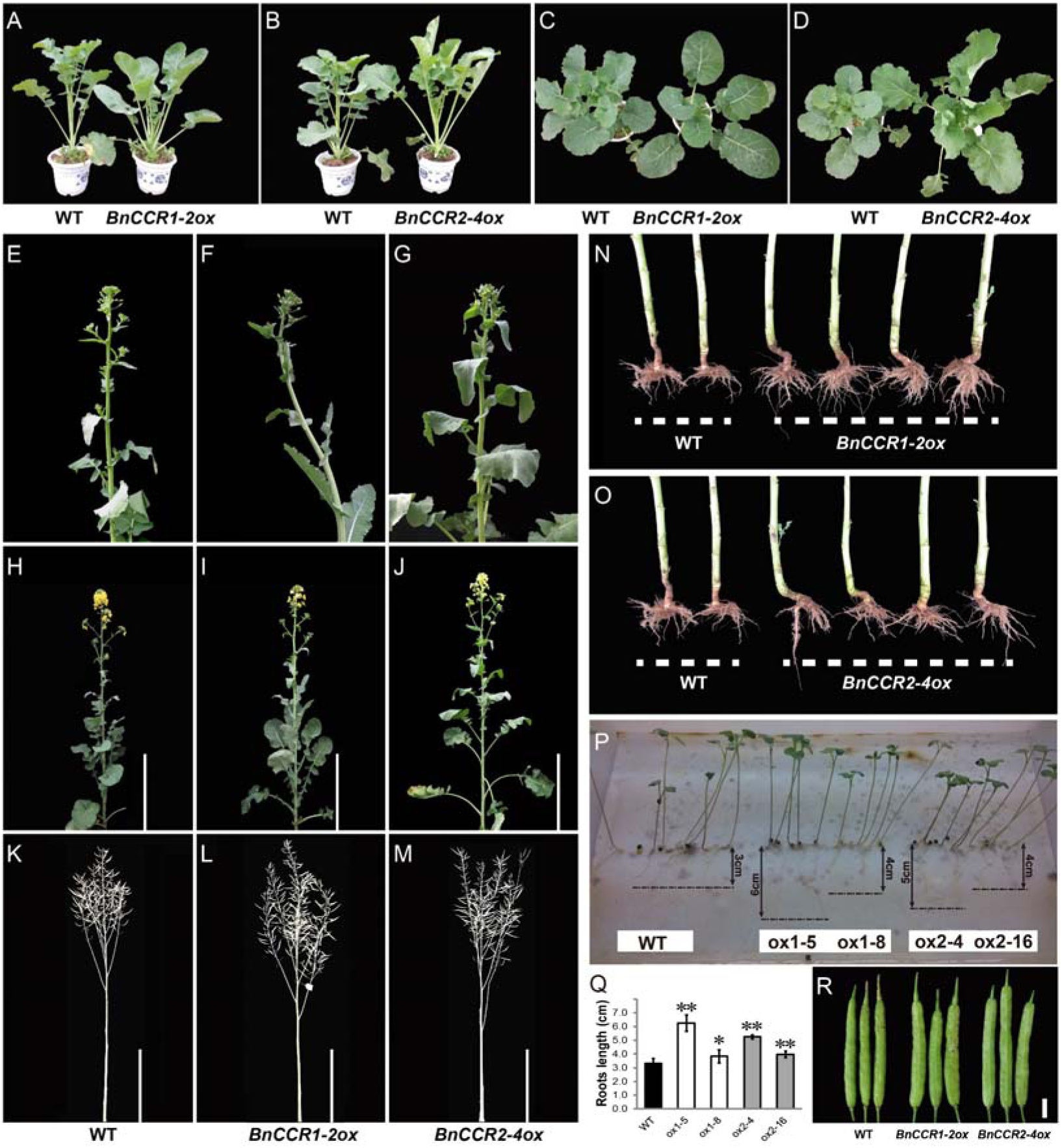
Different phenotypic modifications in *BnCCR1-2ox* and *BnCCR2-4ox* plants. (A-M) Plant phenotypes of different stages. Vegetative stage ([A-D]), middle-later bolting stage ([E-G]), flowering stage ([H-J]), harvest stage ([K-M]). Bending phenomenon of the upper stem of BnCCR1-2ox plants at late bolting stage (F) will disappear after flowering (I). (N, O) root system at mature stage. (P, Q) Root system of one-week old seedlings, and the investigated values with statistic significance. Data represent means ± SD of at least 5 biological replicates. Asterisks indicate that means differed significantly from WT values (*P < 0.05; **P < 0.01, Student’s t tests). (R) Siliques at mature stage, indicating the wider siliques of BnCCR1-2ox and BnCCRR2-4ox compared with WT. Bar = 1cm.

### Yellow-seed traits generated in *BnCCRox* lines

As displayed in Fig. 3A, the seed color of both *BnCCR1-2ox* and *BnCCR2-4ox* turned lighter compared with WT, and *BnCCR1-2ox* showed a stronger effect than *BnCCR2-4ox*. Microscopic investigation of the frozen sections of seed coat (Fig. 3B) and R value obtained through NIRS assay (Fig. 3E) further proved this effect. However, the thickness of the seed coat had no significant difference between *BnCCRox* and WT, which was different from traditionally bred yellow-seeded cultivars which usually had thinner seed coat than black-seeded cultivars (Qu *et al*., 2013; Zhang *et al*., 2013). As expected, the *BnCCR1-2ox* seed coat had an apparent reduction of condensed tannin compared with WT, e.g. ox1-5, ox1-8, ox1-12 and ox1-14 had a decrease of 69%, 29%, 40% and 57% respectively (Fig. 3C and G). But unexpectedly, a significant increase was found for *BnCCR2-4ox* seed coat when compared with WT, e.g. ox2-4, ox2-11, ox2-16 and ox2-25 showed an increase of 88%, 300%, 158% and 83% respectively (Fig. 3C and G), implying looser condensation of the tannin structure caused by unknown mechanisms. Furthermore, all *BnCCRox* lines had a significant reduction of the thousand-seed weight, with *BnCCR1-2ox* lines reduced by 10%-20%, and *BnCCR2-4ox* lines reduced by 15%-30% (Fig. 3D). Surprisingly, the NIRS detection results showed that the total glucosinolates content of *BnCCR1-2ox* lines had a considerable increase compared with WT and a decline in *BnCCR2-4ox* lines (Fig. 3F).

**Fig. 3.**
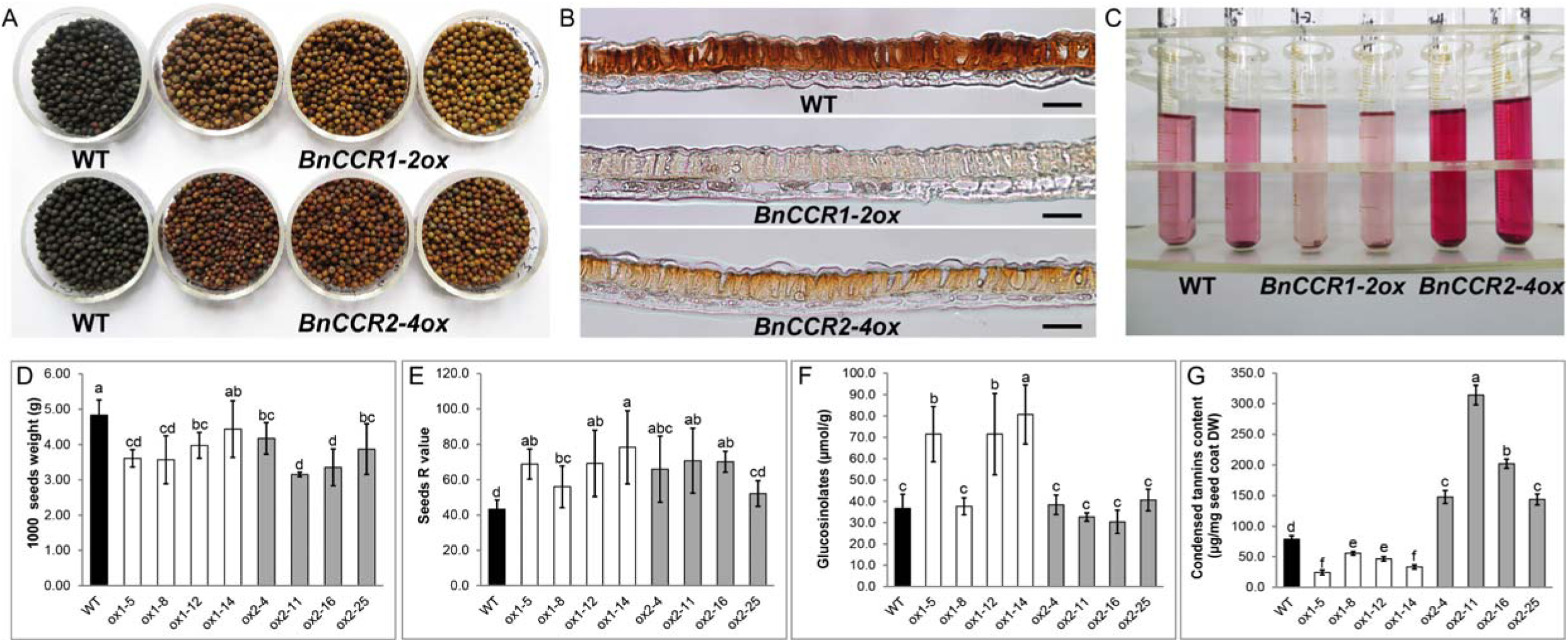
Yellow seeds from T3 plants of *BnCCR1-2ox* and *BnCCR2-4ox* in contrast with black seeds from WT plants. (A, B) seeds and seed coat cross-sections, highlighting the yellow-seed trait caused by seed color lightening by CCR overexpression. Bar = 50µm. (C) extractable insoluble condensed tannins from seed coat, showing deeper color of BnCCR2-4ox as compared with BnCCR1-2ox and WT which implies easier extraction. (D) 1000 seeds weight; (D) Seeds R value (higher value means deeper yellow color of the seeds coat); (F) Glucosinolates content (µmol/g); (G) Insoluble condensed tannins content of the seed coat. Data represent means ± SD of at least three biological replicates. Different letters behind the SD indicate statistically significant differences (one-way ANOVA, P < 0.05, Duncan’s test).

### *BnCCR* over-expression greatly changed lignification phenotypes

The frozen cross sections of stem displayed that both *BnCCR1-2ox* and *BnCCR2-4ox* stems had an changed shape with more concave and convex and a wider xylem parts with deeper histochemical staining (Fig. 4A-C), and some extreme *BnCCR1-2ox* lines appeared an ectopic lignin deposition as shown in Fig. 4C. Observed with higher amplification folds of the sections, *BnCCR1-2ox* showed a better developed xylem part, had more number and more concentrated vessels with deeper brown color (indicating more G-lignin unit) in Mäule staining in which G- and S-type lignin units could be stained brown and red respectively (Fig. 4F and G, early flowering stage), with brighter red phloroglucinol-HCl staining (Fig. 4H, mature stage) and brighter blue fluorescence (Fig. 4I and J, early flowering stage) as compared with WT. These indicated that *BnCCR1-2ox* stems contained higher level of lignin content compared with WT. *BnCCR2-4ox* stem sections also displayed a better developed xylem part and interfascicular fiber part. Although its vessel number and size were not obviously different from WT, there was a significant deeper red color (indicating S-lignin unit) in both Mäule staining (Fig. 4F and G) and phloroglucinol-HCl staining (Fig. 4H), especially to the interfascicular fiber cells wall, and brighter blue florescence under UV light (Fig. 4I and J) compared with WT. This result indicated that *BnCCR2-4ox* stems had higher proportion of S-lignin besides with higher total lignin content. Similar enhancement trends of the lignification pattern were also found in detection results of siliques (Fig. 4K-N).

**Fig. 4.**
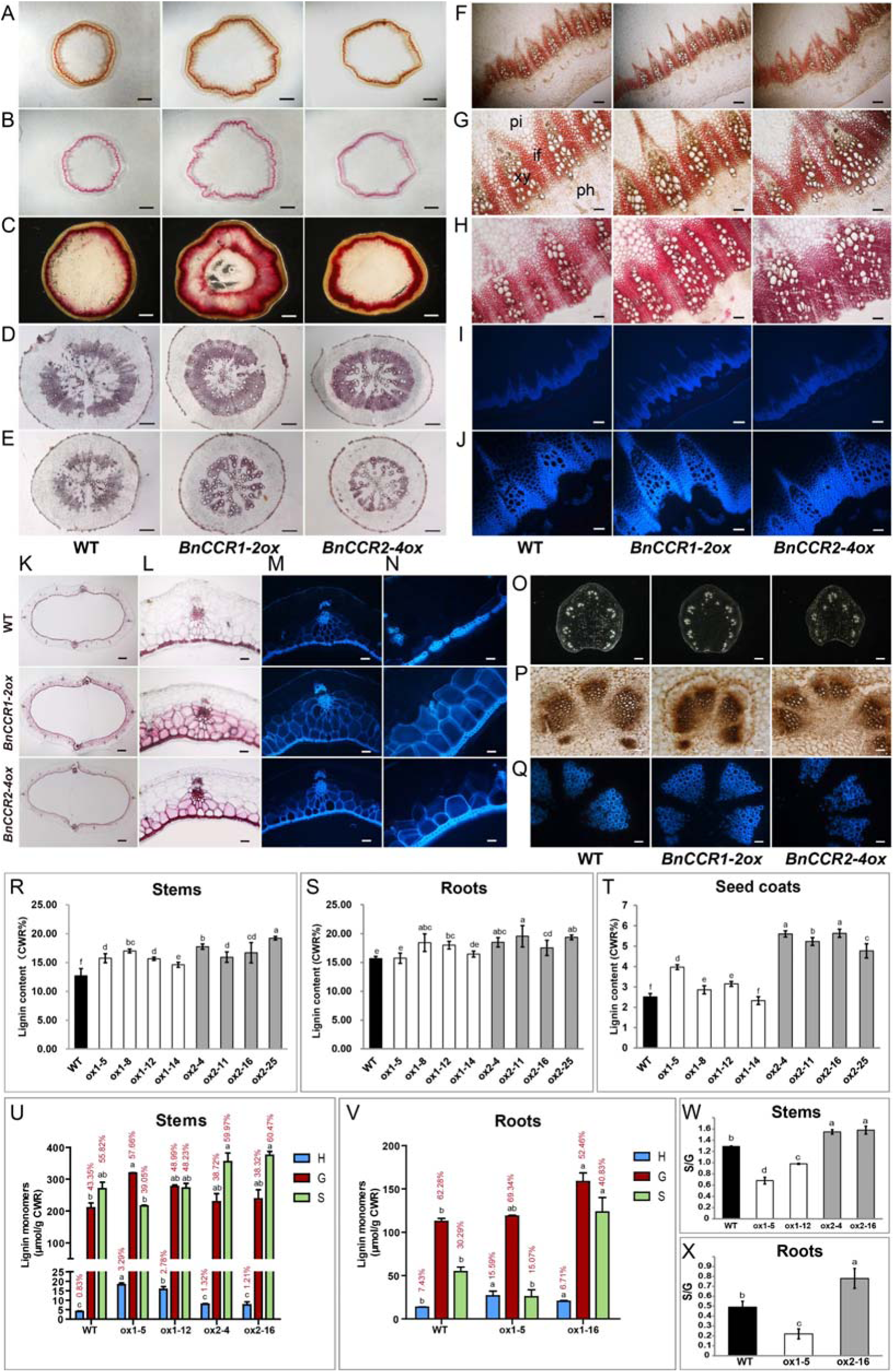
*BnCCR1-2ox* and *BnCCR2-4ox* plants show differently fortified lignification patterns in various organs. (A-C) Whole cross frozen stem-sections at early flowering stage ([A, B]), and whole cross free hand stem-section at harvesting stage (C). (A) Mäule staining; (B, C) phloroglucinol-HCl staining. Bar = 2mm. (D, E) Cross-sections at mature stage of roots with more interfascicular fibers (D) and less interfascicular fibers (E), stained with phloroglucinol-HCl method. Bar = 500 μm. (F-J) Cross-sections of different parts of the stem at different developing stages. Middle-lower part of the stem at early flowering stage using Mäule staining with different amplification folds ([F, G]). Middle-lower part of the stem at mature stage with phloroglucinol-HCl staining (H). Autofluorescence of the middle stem at early flowering stage under UV light with different amplification folds ([I] and [J]). if, interfascicular fiber; ph, phloem; pi, pith; xy, xylem. (F, I), bar = 200 μm; (G), (H, J), bar = 80 μm. (K-N) Silique wall cross-sections at mature stage. Whole sections (K); Local sections ([I-N]). (K-L) Phloroglucinol-HCl staining; (M, N) Autofluorescence viewed under UV light. (K), bar = 500 μm; (L) and (M), bar = μ100 m; (N), bar = μm. (O-Q) Cross sections of the middle petiole with different methods. The whole cross-sections without any treatment (I); The vascular bundle with Mäule staining (P); The autofluorescence of the vascular bundle under UV light (Q); bar = 50 μm. (I), bar = 1000 μm; (P), bar = 100 μm; (Q), bar = 50 μm. (R-T) Total lignin content analysis of stems (r), roots (s) and seed coats (t) was carried out by AcBr method. CWR, cell wall residue. Values are means ± SD of at least three biological replicates. Different letters above the bars indicate statistically significant differences (one-way ANOVA, P < 0.05, Duncan’s test). (U-X) Lignin monomer compositions of stems (U) and roots (V) were measured by thioacidolysis method. S/G ratios in stems and roots are displayed in (W) and (V), respectively. Values are means ± SD of at least three biological replicates. Different letters and values above the bars indicate statistically significant differences (one-way ANOVA, P < 0.05, Tukey’s test) and respective percentage of the corresponding lignin monomers.

This study also reveals that rapeseed has two types of roots: type I with higher proportion of interfascicular fibers, while type II with higher proportion of vessels (Fig. 4D and E). In *BnCCR1-2ox* and *BnCCR2-4ox* lines, both type I and type II roots displayed a better developed xylem tissues with bigger size and more concentrated vessels, and had a deeper staining by phloroglucinol-HCl method compared with WT (Fig. 4D and E). Besides, type II roots of *BnCCR1-2ox* showed a significantly better vessel development (Fig. 4E) than WT and *BnCCR2-4ox*. Through comparison of the above-mentioned histochemical staining assay or fluorescence excitation assay of stems and roots, *BnCCR1-2ox* had better developed vascular bundles, while *BnCCR2-4ox* had better developed interfascicular fibers than WT (Fig. 4D, G, H and J).

For leaf traits, the petiole of *BnCCR1-2ox* tended to be more circular with more and better developed vascular bundles (Fig. 4O and Q). But for *BnCCR2-4ox* lines, it was smaller than WT and developed asymmetrically with fewer and less developed vascular bundles (Fig. 4O and Q), maybe this was one reason that caused *BnCCR2-4ox* leaves much easier to bend and rollover than WT when under strong sunlight and high temperature conditions. Moreover, *BnCCR1-2ox* lines had a deeper brown color in Mäule staining and had a brighter blue fluorescence under UV light than WT (Fig. 4P and Q), which indicated that *BnCCR1-2ox* petiole xylem contained higher lignin content as well as higher G-lignin proportion. The *BnCCR2-4ox* petiole sections displayed no better Mäule staining, and the blue fluorescence was significant weaker than WT (Fig. 4Q), which indicated that *BnCCR2-4ox* petiole xylem contained less lignin content which was consistent with phloroglucinol-HCl staining of leaves (Supplementary Fig. S12J).

### Notable changes in content and structure of lignins in *BnCCR* over-expression lines

Biochemical analysis showed that the acetyl-bromide soluble lignin content of both *BnCCR1-2ox* and *BnCCR2-4ox* was significantly increased compared to WT (Fig. 4R-T). The lignin content of stems of *BnCCR1-2ox* lines ox1-5, ox1-8, ox1-12 and ox1-14 increased by 24%, 34%, 23% and 15%, respectively (Fig. 4R). For *BnCCR2-4ox* lines ox2-4, ox2-11, ox2-16 and ox2-25, their stem lignin content increased by 40%, 25%, 31% and 51%, respectively (Fig. 4R). In roots, the lignin content of ox1-5, ox1-8, ox1-12, ox1-14, ox2-4, ox2-11, ox2-16 and ox2-25 increased by 0.4%, 18%, 15%, 5%, 18%, 25%, 12% and 23% compared with WT, respectively (Fig. 4S). Similar trends also happened in seed coat of *BnCCR1-2ox* lines (Fig. 4T). Similar to tannin detection result, the seed coat lignin content of the *BnCCR2-4ox* lines increased by 92%-130% (Fig. 4T), implying possible decreased lignin polymerization.

S/G ratio, which was typically used to characterize the lignin structure, changed significantly in both stems and roots of *BnCCRox* lines. In the stems of both *BnCCR2-4ox* and WT, their lignin had higher content of S-unit than G-unit, and the S/G ratio was increased by 20.2% and 22.5% in ox2-4 and ox2-16 compared with WT (Fig. 4W). It was mainly attributed to their greater proportion of S-lignin content (Fig. 4U and W; Supplementary Fig. S15), which was in consistent with the Mäule staining results. However, in the case of the stems of *BnCCR1-2ox* lines, the G-unit, other than S-unit, served as the main lignin units, so it was contrary to *BnCCR2-4ox* and WT. Due to increased G-unit proportion, the S/G ratio in the stems of ox1-5 and ox1-12 decreased by 47.3% and 24.0%, respectively, compared to WT (Fig. 4U and W; Supplementary Fig. S15). In roots, the alteration tendency of S/G ratio of ox1-5 and ox2-16 was similar to that in stems, decreased to 0.22 and increased to 0.78, respectively as compared with WT (0.49) (Fig. 4X).

Another striking change was H-lignin proportion, which generally had trace amount in dicotyledonous stems. The H-lignin content of ox1-5 and ox1-12 was about four and three folds of the WT respectively, while was increased by only about 50% in ox2-4 and ox2-16 (Fig. 4U; Supplementary Fig. S15). In the root of ox1-5, the H-lignin content also had an obvious increase, reaching about two folds of WT amount (7.43% of WT VS 15.59% of ox1-5) (Fig. 4V; Supplementary Fig. S15). In conclusion, the lignin structure of *B. rapus* was undoubtedly modified after the manipulation of *BnCCR* genes. However, the NIRS results indicated that cellulose and hemicellulose contents of both *BnCCR1-2ox* and *BnCCR2-4ox* lines had no significant difference when compared with WT (Supplementary Fig. S16).

### Metabolic remodeling of phenylpropanoid and glucosinolate pathways by over-expressing *BnCCR1* and *BnCCR2*

The drastic alteration of seed color, seed coat condensed tannin content, lignin content, lignin structure and plant phenotypes all indicated flux change within and outside the lignin biosynthetic network in *BnCCR* overexpressors.

Conforming to prediction, most phenolic compounds synthesized at the downstream of CCR were apparently increased in the stems of ox1-5 and ox2-16 lines, including sinapoylhexose, sinapic acid, sinapoyl malate, ferulic acid, feruloyl malate, *p*-coumaraldehyde and 1,2-disinapoylglucoside, with an increase of about 1-10 folds, and ox2-16 displayed stronger effects than ox1-5 (Fig. 5A; Supplementary Table S4). However, flavonoids in ox1-5 stems were reduced to 3%-71% of WT, including km-3-O-sophoroside-7-O-glucoside, rutin, is-3-sophoroside-7-glucoside, km-3-O-sinapoylsophoroside-7-O-glucoside, qn-3-O-sophoroside, qn-3-O-glucoside, km-3-O-glucoside and is-3-O-glucoside; and ox2-16 showed the same trend as ox1-5 but with a less extent (Supplementary Table S4). Leaf extracts of ox1-5 and ox2-16 had similar variation tendency as in stems, but some compounds were undetectable in leaves or had an opposite change (Supplementary Table S4). Metabolites profiling was also performed on 30 DAP seeds. As expected, the CCR-downstream compounds sinapic acid and disinapoylgentiobiose were significantly increased in ox1-5 and ox2-16 (Supplementary Table S5). The contents of the most compounds of epicatechin, procyanidin, epicatechin polymers and other important flavonoids were significantly reduced (by even more than 90% for some flavonoids) in ox1-5 compared with wild type, and ox2-16 showed the same trend as ox1-5 but with a less extent (Fig. 5B; Supplementary Table S5).

**Fig. 5.**
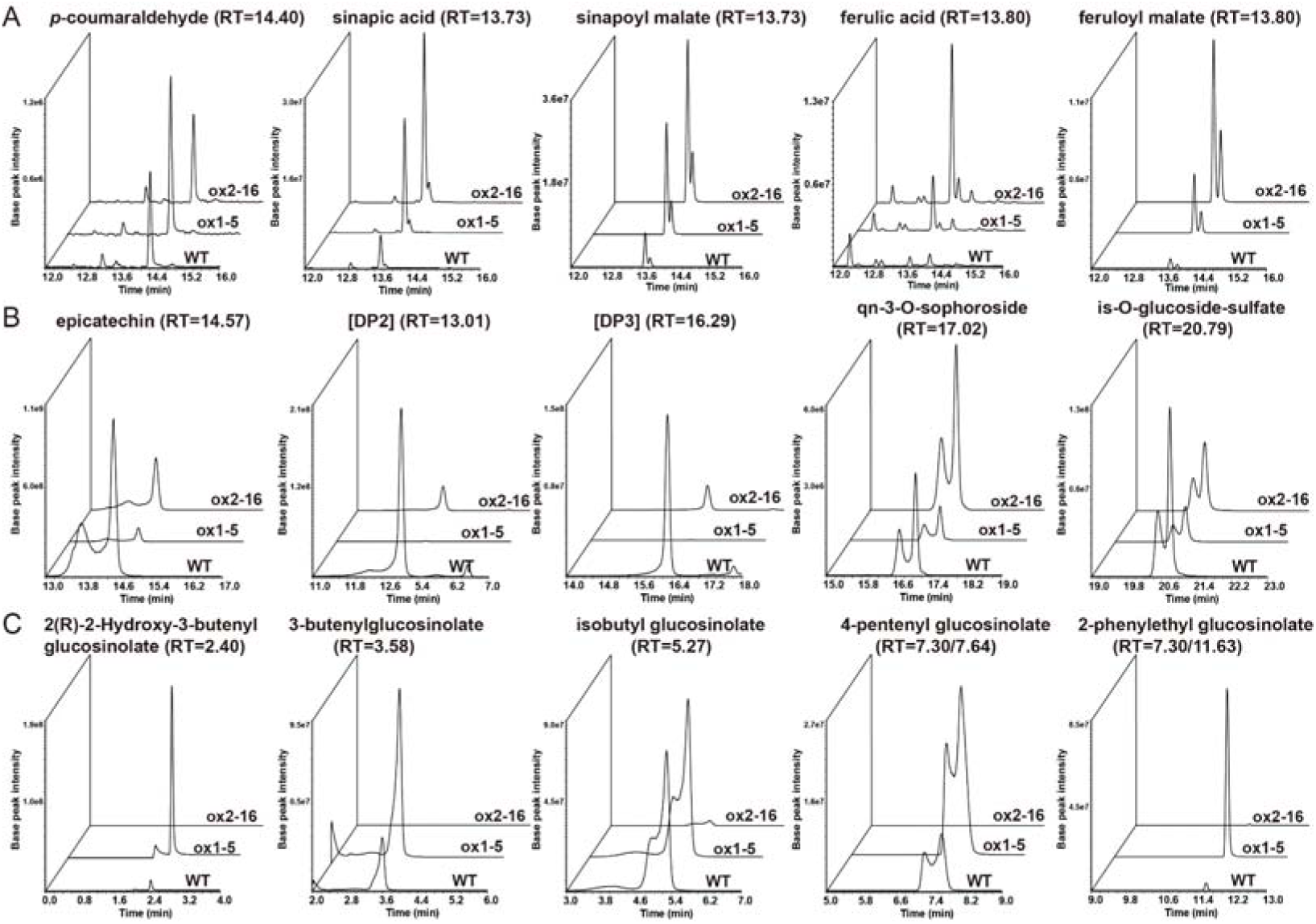
Secondary metabolites are distinctly modified in stems of *BnCCR1-2ox* and *BnCCR2-4ox* plants. Secondary metabolites were determined by UPLC-HESI-MS/MS. (A) Major soluble metabolites related to lignin pathway and its derivative pathways in the stems of ox1-5 and ox2-16 and WT, and see detail information in Supplementary Table S5. (B) Major soluble metabolites related to flavonoid pathway in the seeds of ox1-5 and ox2-16 and WT, and see detail information in Supplementary Table S6. DP, degree of polymerization of the epicatechin unit. (C) Major soluble metabolites related to glucosinolate pathway in the leaves of ox1-5 and ox2-16 and WT, and see detail information in Supplementary Table S5.

In metabolic profiling, glucosinolates were unexpectedly found to be drastically changed by *BnCCR* overexpression. A variety of aliphatic glucosinolates were distinctly differentially deposited between *BnCCRox* lines and WT. For example, 2(R)-2-hydroxy-3-butenyl glucosinolate, 1-S-[(3S)-3-Hydroxy-N-(sulfooxy)-5-hexenimidoyl]-1-thio-beta-D-glucopyranos e, 3-butenylglucosinolate, isobutyl glucosinolate, 4-pentenyl glucosinolate, 5-methylsulfinylpentyl glucosinolate and 5-methylthiopentyl glucosinolate were upregulated to hundreds of folds or even more than 1000 folds in ox1-5 stems as compared with WT, while in ox2-16 stems they were just slightly upregulated or even downregulated (Fig. 5C; Supplementary Table S4). In addition, 4-methylthiobutyl glucosinolate and 6-methylthiohexyl glucosinolate were obviously accumulated in ox1-5 stems (not in ox2-16 and WT stems). According to the quantitative results, glucosinolates had larger amounts of accumulation in leaves than in stems (Supplementary Table S4), and most of them in leaves were several folds higher in ox1-5 than in WT. However, most of them were downregulated by hundreds of folds in the leaves of ox2-16 in comparison with WT. 2(R)-2-Hydroxy-3-butenyl glucosinolate, 1-S-[(3S)-3-Hydroxy-N-(sulfooxy)-5-hexenimidoyl]-1-thio-beta-D-glucopyranos e and 5-methylthiopentyl glucosinolate were even not detectable in the leaves of ox2-16 (Supplementary Table S4). Glucosinolates variation in seed coat was similar to that in the leaf (Supplementary Table S5). Modification of secondary metabolites in petioles was ultimately similar to that in stems (Supplementary Table S6).

### Differential redirection of gene expression in lignin, flavonoid and glucosinolate pathways in *BnCCR1* and *BnCCR2* over-expressors

In *BnCCR1-2ox* lines, *BnCCR1* subfamily itself was significantly upregulated, and *BnCCR2* subfamily showed no significant upregulation. In *BnCCR2-4ox* lines, the *BnCCR2* subfamily itself was undoubtedly greatly upregulated, while *BnCCR1* subfamily was also significantly upregulated (1-3 folds) in the stems, petioles and 30DAP seeds but significantly downregulated by 70% in the leaves (Supplementary Fig. S17). These results indicated that over-expression of *BnCCR1* subfamily had little impact on the expression of *BnCCR2* subfamily, but over-expression of *BnCCR2* subfamily could significantly upregulate or downregulate the expression of *BnCCR1* subfamily depending on different organs.

In the stems of both *BnCCR1-2ox* and *BnCCR2-4ox* lines, the common phenylpropanoid pathway loci *C4H* and *4CL* and the lignin-pathway early-step loci *C3H*, *HCT* and *CCoAOMT* were mildly upregulated (Fig. 6A), and *CCR-*downstream loci *CAD* and *F5H* were slightly upregulated. *COMT* was specific, apparently upregulated in the stems of *BnCCR1-2ox*, but downregulated to less than 10% in the stems of *BnCCR2-4ox* as compared with WT.

**Fig. 6.**
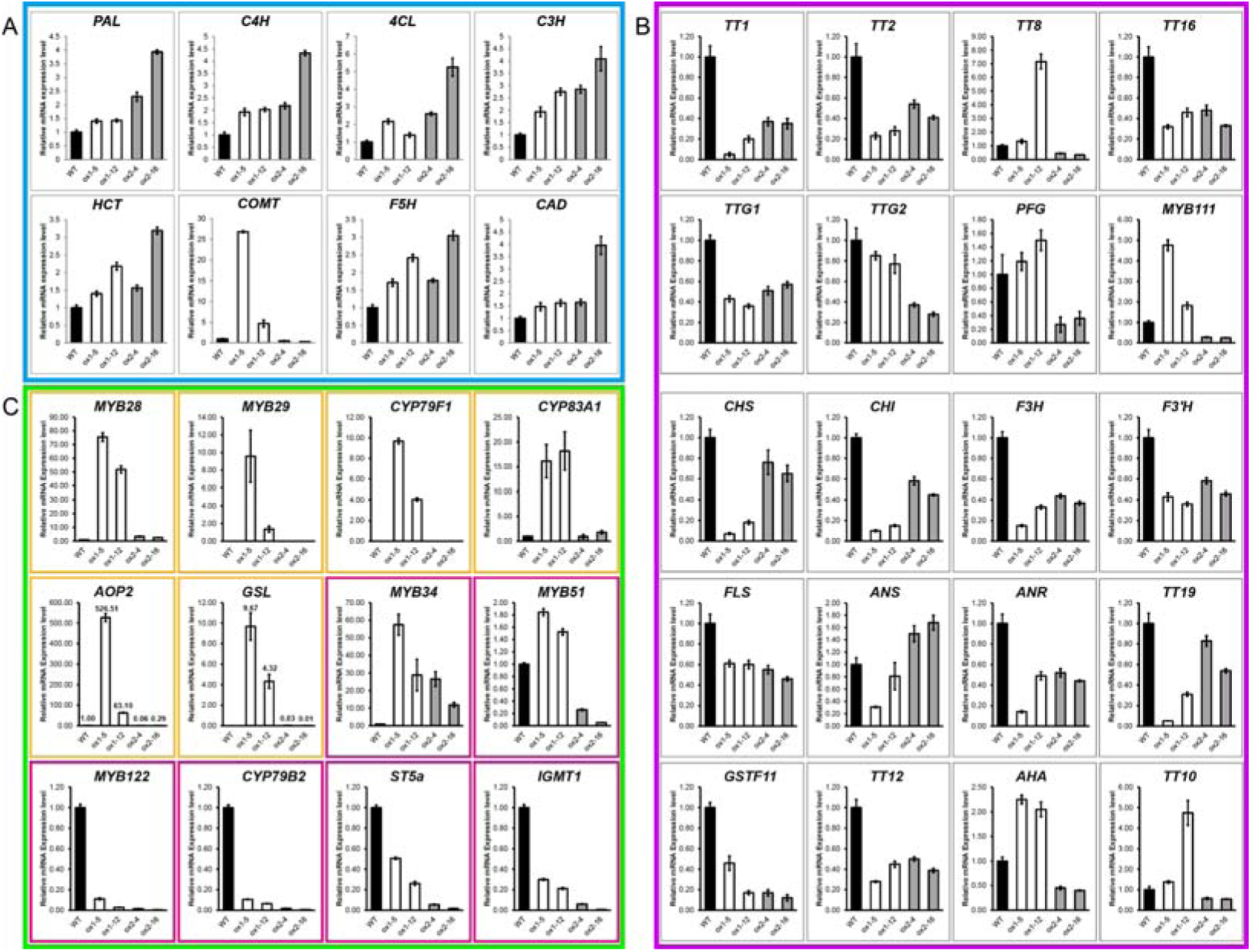
Gene expression patterns of lignin, flavonoid and glucosinolate pathways are tremendously and differentially influenced in *BnCCR1-2ox* and *BnCCR2-4ox* plants. (A) Transcript levels of the genes related to lignin biosynthesis in stems. (B) Transcript levels of the genes related to flavonoid biosynthesis in seeds. (C) Transcript levels of the genes related to glucosinolate biosynthesis in leaves. The MYB29, CYP79F1 and GSL have no expression in WT, hence their expression level in BnCCRox is set relative to zero. The expression levels of other genes are set relative to WT. The yellow frame marks the genes involved in the synthesis of aliphatic glucosinolates. The red frame marks the genes involved in the synthesis of indole glucosinolates.

On the other hand, the expression of the flavonoid biosynthesis pathway was significantly downregulated in *BnCCRox* lines. Regulatory genes *TT1*, *TT2*, *TT16*, *TTG1* and *TTG2*, and structural genes *CHS*, *CHI*, *F3H*, *F3’H*, *FLS*, *ANR*, *TT19*, *GSTF11* and *TT12* were all suppressed to certain extent in 30DAP seeds of both *BnCCR1-2ox* and *BnCCR2-4ox* (Fig. 6B), with less extent in *BnCCR2-4ox* than in *BnCCR1-2ox*. *CHS*, *CHI*, *F3H*, *ANR* and *TT19* were downregulated to less than 20% in the 30DAP seeds of ox1-5 when compared with WT (Fig. 6B). The structural genes *AHA10* and *TT10* and the regulatory genes *TT8* and *MYB111* were significantly downregulated in 30DAP seeds of *BnCCR2-4ox*, but were unexpectedly significantly upregulated in the 30DAP seeds of *BnCCR1-2ox* lines (Fig. 6B). The overall downregulation of the whole flavonoid pathway could account for the reduction of the flavonoids in *BnCCRox* lines.

Most of the genes associated with the aliphatic glucosinolate biosynthesis, such as *MYB28*, *MYB29*, *CYP79F1*, *CYP83A1*, *AOP2* and *GSL*, were significantly upregulated in *BnCCR1ox* lines (Fig. 6C). Especially, the expression of *MYB29*, *CYP79F1* and *GSL* could hardly be detected in WT and *BnCCR2ox* lines, but was considerably expressed in *BnCCR1ox* lines. *AOP2* was a specific locus, which showed considerable downregulation in *BnCCR2ox*, a trend opposite to that in *BnCCR1ox* (Fig. 6C). The expression of most genes involved in indole glucosinolate biosynthesis, such as *MYB122*, *CYP79B2*, *ST5a* and *IGMT1*, was dramatically decreased in both *BnCCR1ox* and *BnCCR2ox* lines, except that *MYB34* was significantly upregulated in both *BnCCR1ox* and *BnCCR2ox* lines (Fig. 6C). *MYB51* was significantly upregulated in *BnCCR1ox* lines but extremely downregulated in *BnCCR2ox* lines, implying that *MYB51* may also be involved in aliphatic glucosinolate biosynthesis in *B. napus*.

## Discussion

### Both *BnCCR1* and *BnCCR2* are crucial for lignification, associated with different monolignols and cellular types

Plants with downregulated CCR activities often displayed reduction in lignin content and alteration in lignin structure depending on different species (Goujon *et al*., 2003; Zhou *et al*., 2010; Prashant *et al*., 2011). Lignin content had a great decrease in *Arabidopsis CCR1* mutant *irx4* (Jones *et al*., 2001) and *CCR*-suppressed transgenic *Arabidopsis* (Goujon *et al*., 2003), tobacco (Piquemal *et al*., 1998), poplar (Leplé *et al*., 2007), *Medicago truncatula* (Zhou *et al*., 2010) and perennial ryegrass (Tu *et al*., 2010).

In this study, anatomical observation and histochemical assays both indicated significant improvement of lignification in stems and roots in both *BnCCR1* and *BnCCR2* over-expressors, and *BnCCR1* over-expressor showed stronger effect on development of vessels and vascular bundles in xylem, while *BnCCR2* over-expressor showed stronger effect on interfascicular fibers.

Furthermore, the S/G ratio was significantly decreased in *BnCCR1-2ox*, mainly due to the stronger increase of G-lignin, the S/G ratio in *BnCCR2-4ox* was significantly increased, mainly attributed to stronger increase of S-lignin. The alteration of S/G ratio caused by *CCR* manipulation always had different trends in different species. A lower S/G ratio, mainly caused by relatively stronger effect of *CCR* downregulation on S-units, was observed on *Arabidopsis irx4* mutants (Patten *et al*., 2005), *CCR*-downregulated poplar (Leplé *et al*., 2007) and *ccr1*-knockout *M. truncatula* mutants (Zhou *et al*., 2010). A higher S/G ratio, mainly caused by relatively stronger effect of *CCR* downregulation on G-units, was found in *CCR*-downregulated tobacco (Chabannes *et al*., 2001), maize (Tamasloukht *et al*., 2011), dallisgrass (Giordano *et al*., 2014) and *CCR2*-knockout *M. truncatula* mutants (Zhou *et al*., 2010). No obvious change of S/G ratio was recorded in poplar (Leplé *et al*., 2007) and perennial ryegrass with *CCR* manipulation (Tu *et al*., 2010).

Additionally, the H-lignin percentage in stems of *BnCCR1-2ox* was 2-3 folds higher than WT, and also more than one-fold higher in roots. Which could be caused by the enhanced accumulation of *p*-coumaraldehyde and expression of *F5H* and *COMT* in *BnCCR1-2* over-expressor. Reasonably, *p*-coumaroyl-CoA and caffeoyl-CoA might serve as the primary substrates for BnCCR1 (as suggested in Fig. 7). In *CCR1*-downregulated perennial ryegrass (Tu *et al*., 2010) and Mu-insertion maize mutant *Zmccr1^-^* (Tamasloukht *et al*., 2011), the H-subunit level was reduced by about 50% and 31% respectively, suggesting that the optimal substrate of CCR1 of perennial ryegrass and maize was *p*-coumaroyl-CoA. On the other hand, BnCCR2 might prefer feruloyl-CoA as its major substrate, because of higher accumulation of S-units and its downstream derivatives sinapate esters, and higher increase in transcript level of *F5H* in *BnCCR2ox* as compared with *BnCCR1ox* (Fig. 7).

**Fig. 7.**
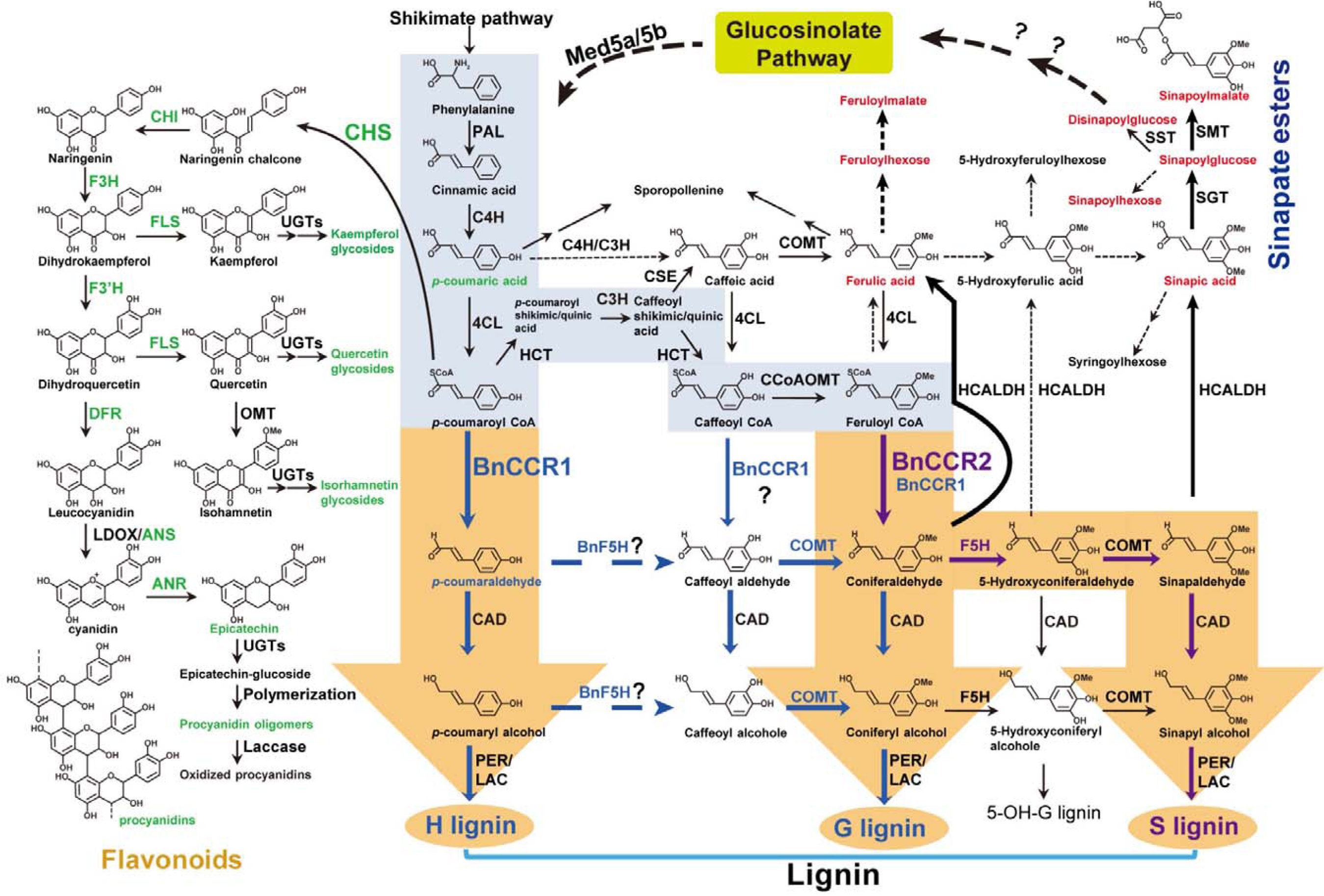
BnCCR1 and BnCCR2 play different roles in phenylpropanoid pathway. The metabolic flux shifts in *BnCCR1-2ox* and *BnCCR2-4ox* plants are displayed in this map. The main route, which is conserved in angiosperms, is marked with the big background arrows (the general phenylpropanoid pathway is marked in light blue, while lignin-specific pathway is marked in brown). The blue arrows and names represent flux enhancement related to *BnCCR1-2ox*. Metabolites accumulation and related gene expression are increased in the stem of *BnCCR1-2ox*, indicating that overexpression of *BnCCR1* subfamily will significantly promote the biosynthesis of G- and H-lignin units. The purple arrows and names represent flux enhancement related to *BnCCR2-4ox*. Metabolites accumulation and related genes expression are increased in the stem of *BnCCR2-4ox*, indicating that overexpression of *BnCCR2* subfamily will significantly promote the biosynthesis of S-lignin unit. The metabolites and the enzymes downregulated in both *BnCCR1-2ox* and *BnCCR2-4ox* plants are marked in green, whereas those with upregulation are in red. Unexpectedly, glucosinolates deposition and pathway gene expression are significantly and differently remodeled in *BnCCR1-2ox* and *BnCCR2-4ox* plants. Dashed arrows represent unknown or unauthenticated routes. Arrows with a question mark are suggested pathways in this study. Two successive arrows represent two or more metabolic conversions. The enzymes and their abbreviations are as follows: ANR, anthocyanidin reductase; ANS, anthocyanidin synthase; CAD, cinnamyl alcohol dehydrogenase; 4CL, 4-coumarate:CoA ligase; C3H, *p*-coumarate 3-hydroxylase; C4H, cinnamate 4-hydroxylase; CCoAOMT, caffeoyl-CoA O-methyltransferase; CCR, cinnamoyl-CoA reductase; CHI, chalcone isomerase; CHS, chalcone synthase; COMT, caffeic acid *O*-methyltransferase; CSE, caffeoyl shikimate esterase; DFR, dihydroflavonol 4-reductase; F3H, flavanone 3-hydroxylase; F3’H, flavonoid 3’-hydroxylase; F5H, ferulate 5-hydroxylase; FLS, flavonol synthase; HCALDH, hydroxycinnamaldehyde dehydrogenase; HCT, hydroxycinnamoyl-CoA:shikimate/quinic hydroxycinnamoyl transferase; LAC, laccase; LDOX, leucoanthocyanidin dioxygenase; Med: mediator; PAL, phenylalanine ammonia-lyase; PER, peroxidase; SGT, sinapate 1-glucosyltransferase; SMT, sinapoylglucose:malate sinapoyltransferase; SST, sinapoylglucose:sinapoylglucose sinapoylglucosetransferase; UGT, uridine diphosphate glycosyltransferase.

### Upregulation of *BnCCR* genes enhanced lodging resistance, modified morphology, but did not improve disease and UV-B resistance

The expression perturbation of the genes located at the lignin biosynthetic pathway was often accompanied with defects in plant growth and development depending on which gene was targeted. For the severely silenced *CCR* plants, phenotypic abnormalities with irregular vessels usually arise, including plant size reduction, delayed flowering, delayed senescence, retarded seed development, biomass yield reduction and compromised pathogen defense (Leplé *et al*., 2007; Zhou *et al*., 2010; Vanholme *et al*., 2012; Van Acker *et al*., 2014; Xue *et al*., 2015; De Meester *et al*., 2018; De Meester *et al*., 2020).

Firstly, *BnCCR* over-expression improved lodging resistance compared with WT, especially for *BnCCR1-2ox* (Fig. 2; Supplementary Fig. S12). When *CCR1* was reintroduced into *A. thaliana ccr1* mutants under the control of the ProSNBE promoter, specific expression of *CCR1* in the protoxylem and metaxylem vessel cells showed a full recovery in vascular integrity (De Meester *et al*., 2018). The breaking-resistance of both *BnCCR1-2ox* and *BnCCR2-4ox* plants was greatly enhanced in comparison with WT, with larger effect in *BnCCR1-2ox* lines than in *BnCCR2-4ox* lines. These results imply an improved lodging resistance of *BnCCRox* lines due to better development and growth in both morphology and lignification of roots and stems, which provides significant potential in molecular breeding of rapeseed with enhanced lodging resistance through over-expressing *BnCCR* genes.

Secondly, *BnCCR* over-expression also modified leaf morphology. Our Mäule staining results indicated that rapeseed petiole xylem was exclusively composed of vascular bundles without interfascicular fibers, which could account for better development of leaf veins in *BnCCR1-2ox* plants, since vascular bundles contained higher proportion of G-lignin and *BnCCR1-2* was mainly involved in the biosynthesis of G- and H-lignins. In *Arabidopsis*, *CCR1* also mediated cell proliferation exit for leaf development, and *ccr1-4* mutant had a significantly reduced leaf and plant size compared with WT (Xue *et al*., 2015). However, *BnCCR2-4ox* plants showed less developed leaf vascular bundles (Fig. 4O; Supplementary Fig. S12J), which could possibly be resulted from two reasons: weakened G-unit synthesis (since *BnCCR2-4* overexpression mainly forced S-unit synthesis), and downregulated *BnCCR1* expression in leaves of *BnCCR2-4ox* plants (Supplementary Fig. S17B). Enhanced levels of ABA might also play a role in the alteration of leaf phenotypes of *BnCCRox* plants (Supplementary Fig. S13), as ABA plays important roles in plant growth and development (Cutler *et al*., 2010).

Thirdly, resistance of *BnCCRox* plants to *S. sclerotiorum* and UV-B was not improved, which was in confliction with traditional notion and our original expectation. There are many factors involved in plant disease resistance, such as the epidermal cuticle-wax layers, trichomes, lignified cell walls, phenolics, phytoalexins and pathogenesis-related proteins (Zhao *et al*., 2007a). In the present study, although the lignin content was increased through *BnCCR* over-expression, the other important factors correlated to the plant resistance were not enhanced or even suppressed by flux competition between lignin pathway and flavonoid and other pathways, which could be reflected by the changes of flavonoid and glucosinolate metabolic profiles. Besides, decrease of epidermal wax layer was observed on leaves of *BnCCR1-2ox* plants compared with *BnCCR2-4ox* and WT (Supplementary Fig. S18).

### *BnCCR1* and *BnCCR2* differentially affect plant development progress

During the progress of the growth and development of *BnCCRox* plants, the upper stem of *BnCCR1-2ox* plants at bolting stage was more easily to bend, but *BnCCR2-4ox* plants had no such case (Fig. 2F; Supplementary Fig. S12A). As reported, S-lignin percentage in stems would gradually increase during plant growth process (Tu *et al*., 2010; Giordano *et al*., 2014). Mäule staining of stem cross sections in this study revealed similar trend (Supplementary Fig. S19), suggesting that S-lignin might play an important role for maturation and mechanical strength of stems. (Kaur *et al*., 2012) found that the stem of *CAD*-downregulated *Nicotiana* plants (ir-CAD) presented a rubbery phenomenon, and the S/G ratio was significantly reduced. Here the S/G ratio was also significantly decreased in *BnCCR1-2ox* plants (1.29 in WT, 0.68 in ox1-5, and 0.98 in ox1-12). Moreover, our results showed that in WT the S/G ratio in stem (1.29) was significantly higher than in root (0.49), which might be one factor contributing to the more flexible texture of roots compared with stems, although total lignin content was higher in root than in stem (Fig. 4R and S). Lower percentage of S-units might contribute to a lower level of stiffness of the stem, together with the lower lignin content in the upper stem, which might be one reason of the bending phenomenon of *BnCCR1-2ox* at bolting stage. However, when *BnCCR1-2ox* plants entered flowering stage, the bending phenomenon disappeared, which might be due to increase in total lignin content and S-lignin percentage in stems (Supplementary Fig. S19). Relationship between S/G ratio and the texture of the plant stems was summarized in the Supplementary Table S7. These results suggest that higher S/G ratio might contributes to stronger stiffness in plant stems, which is manifested during reproductive growth.

In this study, *BnCCR1-2ox* plants flowered distinctly later than WT and *BnCCR2-4ox* plants, implying that the function of Brassicaceae *CCR1* is also associated with plant development progress. There was an evolutionarily conserved mechanism between cell wall biosynthesis and production of flowers (Vermerris *et al*., 2002). The delayed accumulation of S-lignin and lowered stem stiffness at bolting stage were reflections of delayed development progress and flowering in *BnCCR1-2ox* plants.

### *BnCCR1* and *BnCCR2* exert different flux control on flavonoid pathway and lignin-derivative pathways in *B. napus*

In metabolic engineering, products, precursor steps as well as associated neighbor pathways would be affected via metabolic flux redirection. In lignin engineering, for example, significantly higher amounts of vanillin, ferulic acid, *p*-coumaric acid, coniferaldehyde and syringaldehyde were released from the cell wall samples of *MtCAD1* mutants than the wild-type (Zhao *et al*., 2013). In the *C3’H* (Abdulrazzak *et al*., 2006) or *HCT* (Besseau *et al*., 2007) defect *Arabidopsis*, *C3H1-*downregulated maize (Fornalé *et al*., 2015)*, CCR*-silenced tomato (van der Rest *et al*., 2006) and perennial ryegrass (Tu *et al*., 2010), the accumulation of flavonoids was significantly increased.

Besides enhancement of lignin monomers, our results also showed that both *BnCCR1-2* and *BnCCR2-4* over-expressors had significantly increased levels of various CCR-downstream pathway products when compared with WT, especially for sinapate esters (Fig. 5; Supplementary Tables S4-S6), which could be caused by the enhanced expression levels of corresponding genes *CCR*, *F5H*, *COMT* and *CAD*. Moreover, *BnCCR2-4ox* plants had a stronger effect on the accumulation of sinapate esters in stems than *BnCCR1-2ox* plants, implying that the metabolic route related to BnCCR2 catabolism (mainly S-units synthesis) might be closer to sinapate ester pathway than BnCCR1 (Fig. 5A; Fig. 7).

More strikingly, many compounds synthesized in flavonoid pathway were significantly reduced in *BnCCRox* plants compared with WT. The dramatic reduction in flavonoids was theoretically caused by the reduced availability of *p*-coumaroyl-CoA precursor for CHS as *BnCCR* overexpression attracted more of this common precursor into lignin pathway (Fig. 7). This kind of flux shift could be further evidenced on molecular level (Fig. 6). Seed color degree, metabolic profile and gene expression profile all indicated that suppression impact on flavonoid pathway was greater in *BnCCR1-2ox* plants than in *BnCCR2-4ox* plants (Figs 3 and 6; Supplementary Tables S4-S6). Which could result from the fact that the metabolic routes of BnCCR1 catabolism (mainly H- and G-units synthesis) were closer to flavonoid pathway than that of BnCCR2 (mainly S-unit synthesis) as suggested in Fig. 7.

### *BnCCR1* and *BnCCR2* play different roles in crosstalk between phenylpropanoid pathway and glucosinolate pathway

There were a few reports on crosstalk effect of glucosinolate pathway on phenylpropanoid pathway (Hemm *et al*., 2003; Kim *et al*., 2015; Kim *et al*., 2020). In *Arabidopsis*, the accumulation of phenylpropanoids were significantly suppressed in *ref5-1* mutant, and *REF5* was proved to encode CYP83B1 which was involved in biosynthesis of indole glucosinolates (Kim *et al*., 2015). Defect of phenylpropanoids deposition was also detected in *Arabidopsis ref2* mutant, as *REF2* encoded CYP83A1 which played a role in aliphatic glucosinolate pathway (Hemm *et al*., 2003). In low-lignin *c4h*, *4cl1*, *ccoaomt1* and *ccr1* mutants of *Arabidopsis*, transcripts of some glucosinolate biosynthesis genes were more abundant (Vanholme *et al*., 2012). However, to date there is no systemic study on crosstalk effect of phenylpropanoid pathway on glucosinolate pathway.

In present study, the aliphatic glucosinolates were significantly increased in *BnCCR1-2ox* (Supplementary Table S4), which could result from upregulated expression of *MYB28*, *MYB29*, *CYP79F1*, *CYP83A1*, *AOP2* and *GSL* in the leaves of transgenic plants (Fig. 6). Among them, the expression of *MYB29*, *CYP79F1* and *GSL* could almost only be detected in *BnCCR1-2ox*, not in WT. Reasonably, the undetectable *MYB29* expression and extremely lower expression of *MYB51*, *MYB122*, *AOP2*, *CYP79B2*, *ST5a* and *IGMT1* in *BnCCR2-4ox* plants could be responsible for the decrease of glucosinolate content. Because the WT material ZS10 of this study was a “double-low” (low erucic acid, low glucosinolates) commercial cultivar, the low proportion of indole and aromatic glucosinolates in WT and the limited change of indole and aromatic glucosinolates in *BnCCR*-overexpressing plants might be caused by breeding impairment of respective biosynthesis pathways of these two types of glucosinolates. If a “double-high” stock was used as WT for *BnCCR* transformation, the effect on indole and aromatic glucosinolates deposition might not be so mild. Taken together, both *BnCCR1-2* and *BnCCR2-4* distinctly affect glucosinolate biosynthesis, but have divergent or almost opposite effects.

Crosstalk effect of glucosinolate pathway on phenylpropanoid pathway could be linked through PAL degradation that mediated by Med5-KFBs-dependent manner (Kim *et al*., 2020). However, what mechanism is involved in change of glucosinolate biosynthesis by manipulating phenylpropanoid genes, such as *CCR*, needs to be clarified. Our qRT-PCR results indicate that at least transcription regulation is involved, but whether mediator-mediated protein degradation in glucosinolate pathway is also involved deserves future study.

## Supplementary data

**Fig. S1.** Southern blot detection of *CCR* subfamily genes in *B. napus*, *B. rapa* and *B. oleracea*.

**Fig. S2.** Multi-alignment indicates that *Brassica* CCRs contain complete structural features as those in model plants *A. thaliana* and popular.

**Fig. S3.** Overall expression patterns of *CCR1*-subfamily and *CCR2*-subfamily show distinct organ-specificity and difference between black and yellow seeds in *B. napus*, *B. rapa* and *B. oleracea*.

**Fig. S4.** Expression patterns of *CCR1*-genes and *CCR2*-genes show distinct organ-specificity and difference between black and yellow seeds in *B. napus*.

**Fig. S5.** Expression patterns of *CCR1*-genes and *CCR2*-genes show distinct organ-specificity and difference between black and yellow seeds in *B. rapa*.

**Fig. S6.** Expression patterns of *CCR1*-genes and *CCR2*-genes show distinct organ-specificity and difference between black and yellow seeds in *B. oleracea*.

**Fig. S7.** Overall expression of *BnCCR1*-subfamily and *BnCCR2*-subfamily distinctly respond to various stresses in *B. napus* seedlings.

**Fig. S8.** *BnCCR1*-genes and *BnCCR2*-genes distinctly respond to various stresses in *B. napus* seedlings.

**Fig. S9.** Basta-resistance, GUS-staining and PCR determination of transgenic lines.

**Fig. S10.** qRT-PCR shows distinct upregulation of the target genes in respective T2 lines of *BnCCR1-2ox* and *BnCCR2-4ox*.

**Fig. S11.** Multiple agronomic traits are significantly modified in *BnCCR1-2ox* and *BnCCR2-4ox* lines.

**Fig. S12.** *BnCCR1-2ox* and *BnCCR2-4ox* lines show different growth behaviors and leaf vein strengths.

**Fig. S13.** ABA content is upregulated in leaves of *BnCCR1-2ox* and *BnCCR2-4ox* lines.

**Fig. S14.** Plants of both *BnCCR1-2ox* and *BnCCR2-4ox* do not show enhanced resistance to *S. sclerotiorum* and UV-B.

**Fig. S15.** GC-MS chromatograms show different changes of monolignol proportions in *BnCCR1-2ox* and *BnCCR2-4ox* lines.

**Fig. S16.** Contents of cellulose and hemicellulose in the stems of *BnCCR1-2ox* and *BnCCR2-4ox* lines are not changed based on NIRS detection.

**Fig. S17.** *BnCCR1* expression is obviously altered in *BnCCR2-4ox* lines, whereas *BnCCR2* expression has little change in *BnCCR1-2ox* lines.

**Fig. S18.** Leaf surfaces of *BnCCR1-2ox* and *BnCCR2-4ox* plants are less flat with decreased wax deposition than WT.

**Fig. S19.** Mäule staining of stem sections indicates a trend of increase of S-type lignins during developmental process in *B. napus*.

**Table S1.** Identity parameters of *CCR1*-subfamily and *CCR2*-subfamily genes from *B. napus* and its parental species *B. oleracea* and *B. rapa*.

**Table S2.** cDNA and gDNA basic parameters of *Brassica CCR1* and *CCR2* subfamily genes cloned in this study.

**Table S3.** Basic parameters and important features of *Brassica* CCR1 and CCR2 proteins.

**Table S4.** Contents of major soluble secondary metabolites differentially accumulated in stems and leaves of ox1-5 and ox2-16 compared with WT as revealed by UPLC-HESI-MS/MS.

**Table S5.** Contents of major soluble secondary metabolites differentially accumulated in 30DAP seeds of ox1-5 and ox2-16 compared with WT as revealed by UPLC-HESI-MS/MS.

**Table S6.** Contents of major soluble secondary metabolites differentially accumulated in petioles of ox1-5 and ox2-16 compared with WT as revealed by UPLC-HESI-MS/MS.

**Table S7.** S/G ratio of stem lignin, stem stiffness, and plant phenotype of different angiosperm species.

**Table S8.** Primers used in this study.

## Acknowledgements

We thank Prof Xuekuan Zhang for providing Zhongshuang 10 seeds, Prof Wei Qian for providing the *S. sclerotiorum* source strain, Prof Ningjia He, Dr. Guangyu Ding and Dr. Shiyao Liu for assistance on GC-MS measurements. This research was supported by National Natural Science Foundation of China (31871549, 32001579, 31830067 and 31171177), National Key R&D Program of China (2016YFD0100506), Special financial aid to post-doctor research fellow of Chongqing (XmT2018057), “111” Project (B12006), and Young Eagles Program of Chongqing Municipal Commission of Education (CY200215).

## Author contributions

Y.C., J.L. and N.Y. designed the study; N.Y., B.L., X.L. and Y.X. performed the experiments; Y.C. did molecular and bioinformatic analysis, N.Y., J.L., M.C., K.L., L.W. and Y.L. analyzed or evaluated the results; J.L. and R.W. provided some *Brassica* materials. N.Y. and Y.C. performed final analysis and wrote the article.

## Data availability statement

The data supporting the findings of this study are available from the corresponding author (Jiana Li and Yourong Chai) upon request.

