## Supplementary material for "Two Types of Cinnamoyl-CoA Reductase Function Divergently in Tissue Lignification, Phenylpropanoids Flux Control, and Inter-pathway Cross-talk with Glucosinolates as Revealed in *Brassica napus*": Supplymentary Figures-20210413

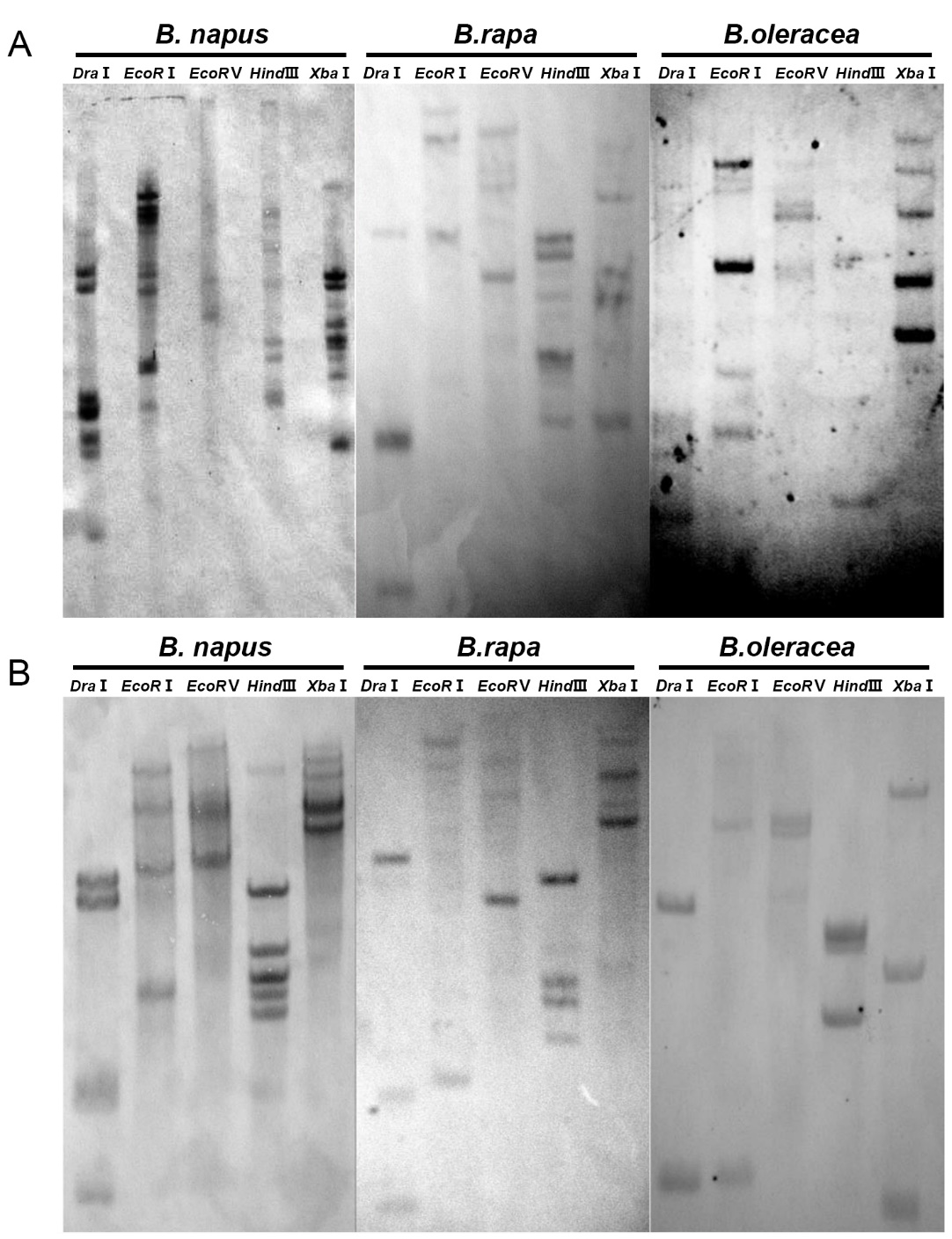

**Fig. S1.** Southern blot detection of *CCR* subfamily genes in *B. napus*, *B. rapa* and *B. oleracea*.

(A) *CCR1* subfamily genes detection.

(B) *CCR2* subfamily genes detection.

The total genomic DNA was fully digested with *Dra*I, *Eco*RI, *Eco*RV, *Hin*dIII and *Xba*I, respectively.

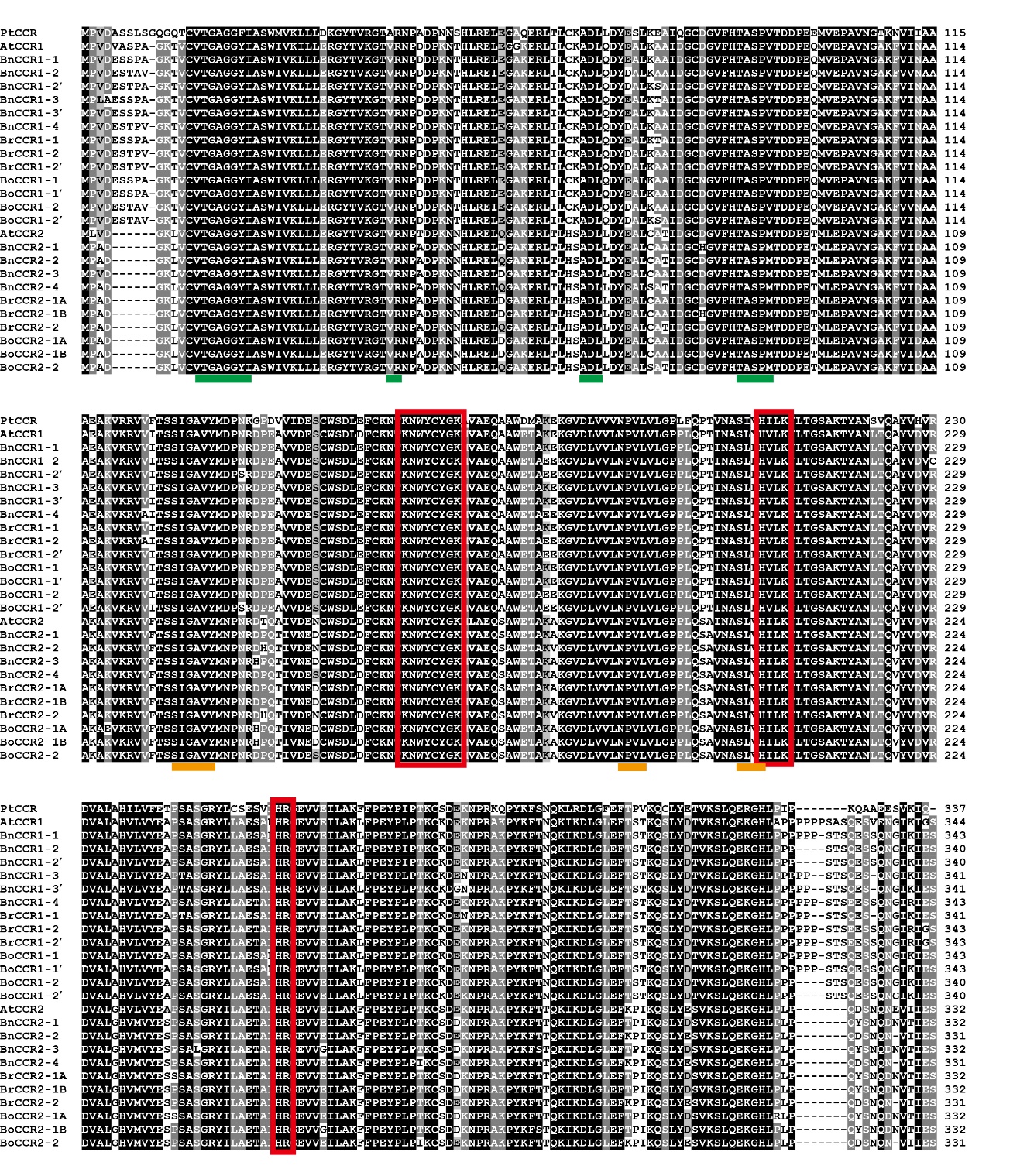

**Fig. S2.** Multi-alignment indicates that *Brassica* CCRs contain complete structural features as those in model plants *A. thaliana* and popular.

The accession numbers of *Brassica* and *A. thaliana* CCRs are the same as in Fig. 1, while the accession number of *Populus tremuloides* CCR (PtCCR) is AAF43141.1. Green bars, NADP binding motifs; Yellow bars, substrate binding motifs; Red boxes, conserved catalytic activity motifs of CCR.

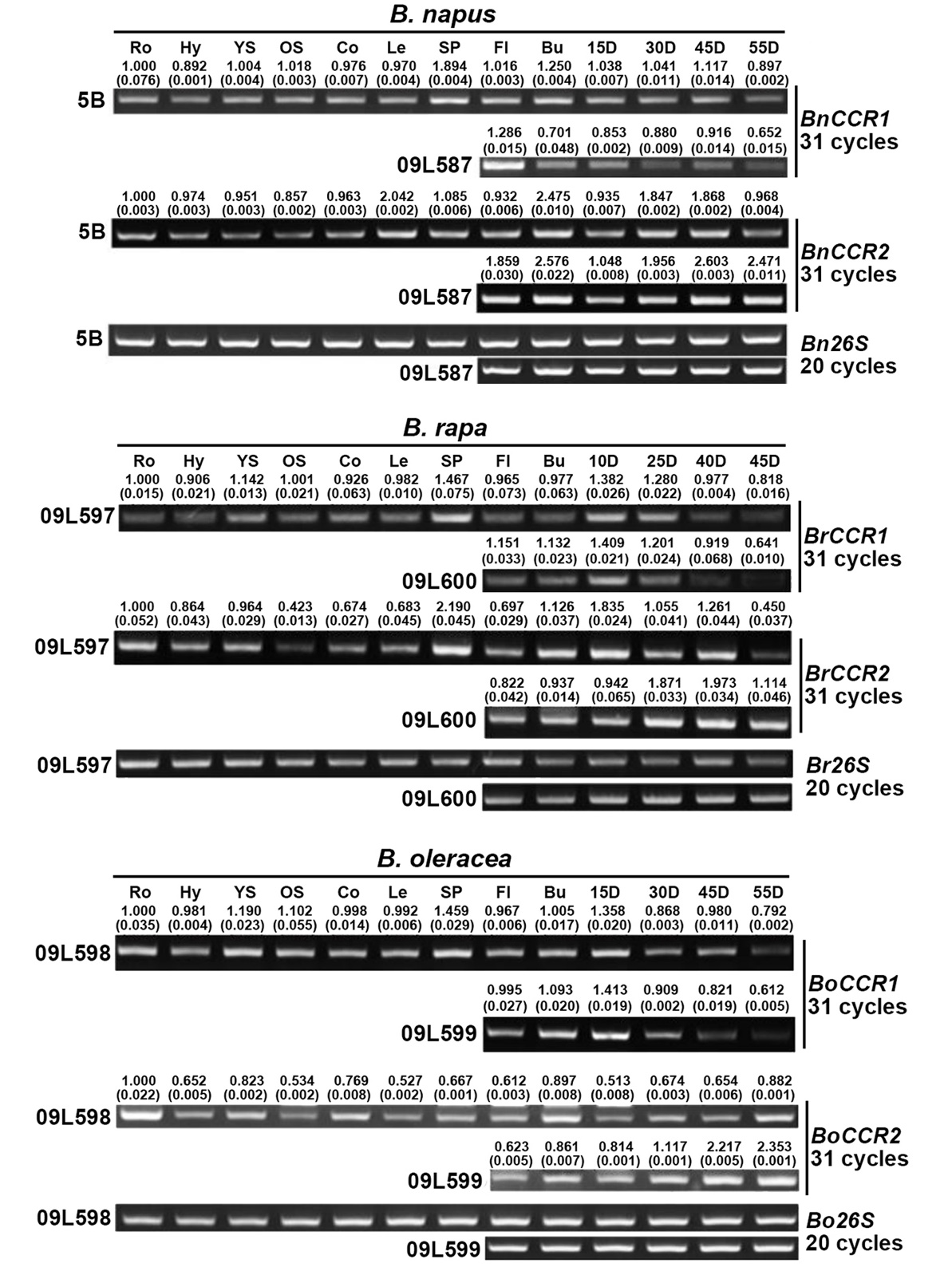

**Fig. S3.** Overall expression patterns of *CCR1*-subfamily and *CCR2*-subfamily show distinct organ-specificity and difference between black and yellow seeds in *B. napus*, *B. rapa* and *B. oleracea*.

5B, 09L597 and 09L598 are black-seed stocks, while 09L587, 09L600 and 09L599 are yellow-seed stocks, of *B. napus*, *B. rapa* and *B. oleracea*, respectively. The value (SEM in brackets) above the sRT-PCR band represents the relative expression level of qRT-PCR result. The expression level in root is set as 1.000 for quantification of other organs. Ro, roots; Hy, Hypocotyl; YS, young stems; OS, old stems; Co, cotyledons; Le, leaves; SP, silique pericarp; Fl, flowers; Bu, buds; 15D-55D, seeds of the corresponding days after pollination.

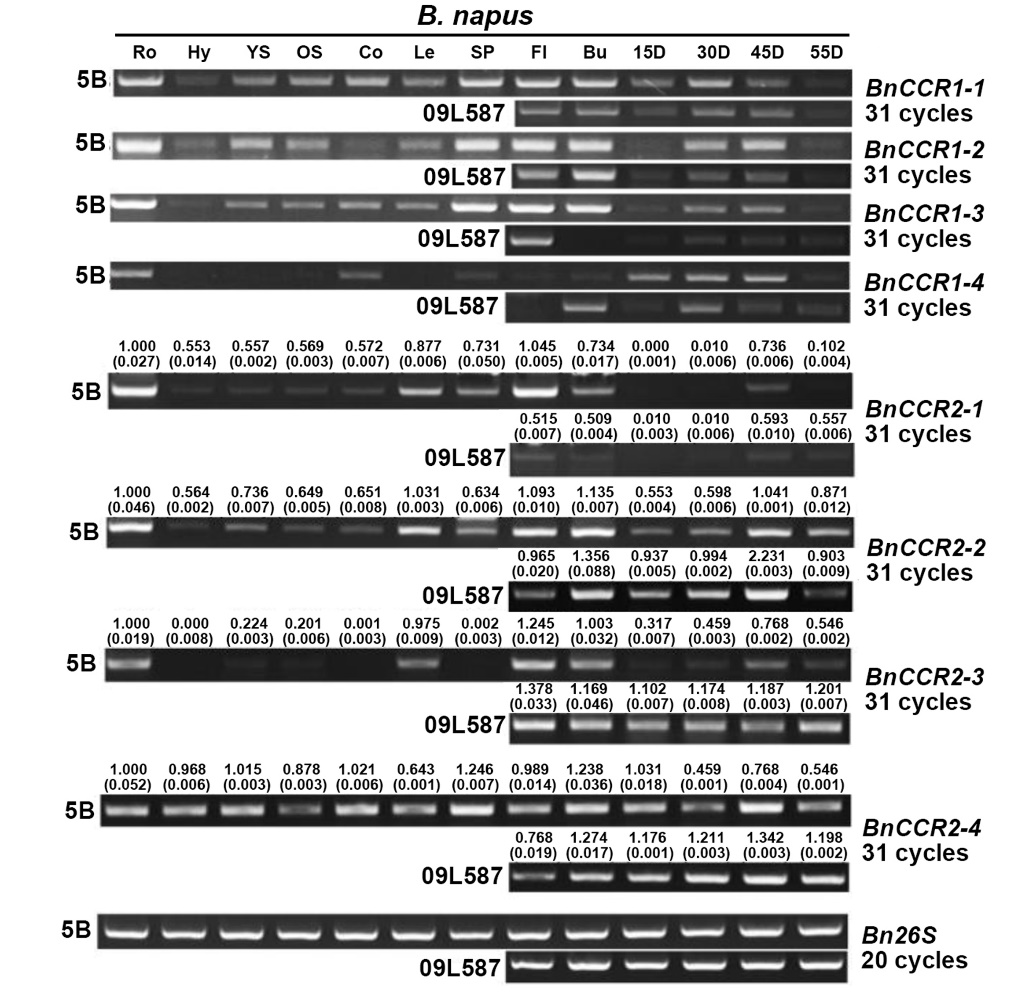

**Fig. S4.** Expression patterns of *CCR1*-genes and *CCR2*-genes show distinct organ-specificity and difference between black and yellow seeds in *B. napus*.

5B and 09L587 are black- and yellow-seed stocks, respectively. The value (SEM in brackets) above the sRT-PCR band represents the relative expression level of qRT-PCR result. The expression level in root is set as 1.000 for quantification of other organs. Ro, roots; Hy, Hypocotyl; YS, young stems; OS, old stems; Co, cotyledons; Le, leaves; SP, silique pericarp; Fl, flowers; Bu, buds; 15D-55D, seeds of the corresponding days after pollination.

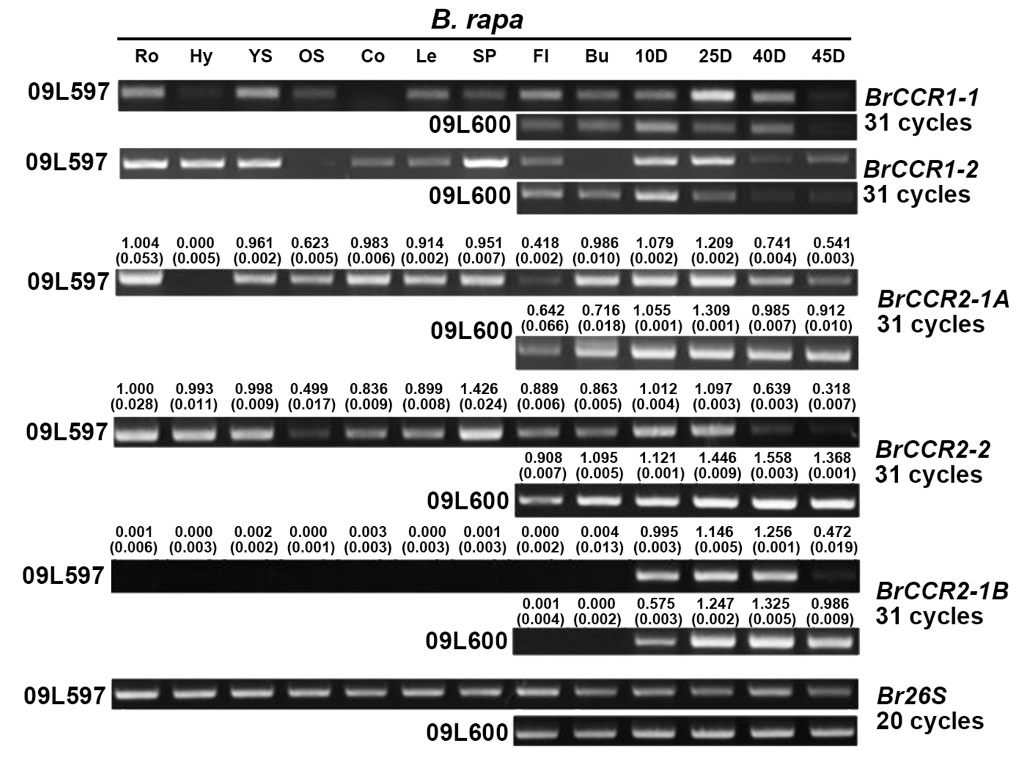

**Fig. S5.** Expression patterns of *CCR1*-genes and *CCR2*-genes show distinct organ-specificity and difference between black and yellow seeds in *B. rapa*.

09L597 and 09L600 are black- and yellow-seed stocks, respectively. The value (SEM in brackets) above the sRT-PCR band represents the relative expression level of qRT-PCR result. The expression level in root of 09L597 is set as 1.000 for quantification of other organs. Ro, roots; Hy, Hypocotyl; YS, young stems; OS, old stems; Co, cotyledons; Le, leaves; SP, silique pericarp; Fl, flowers; Bu, buds; 15D-55D, seeds of the corresponding days after pollination.

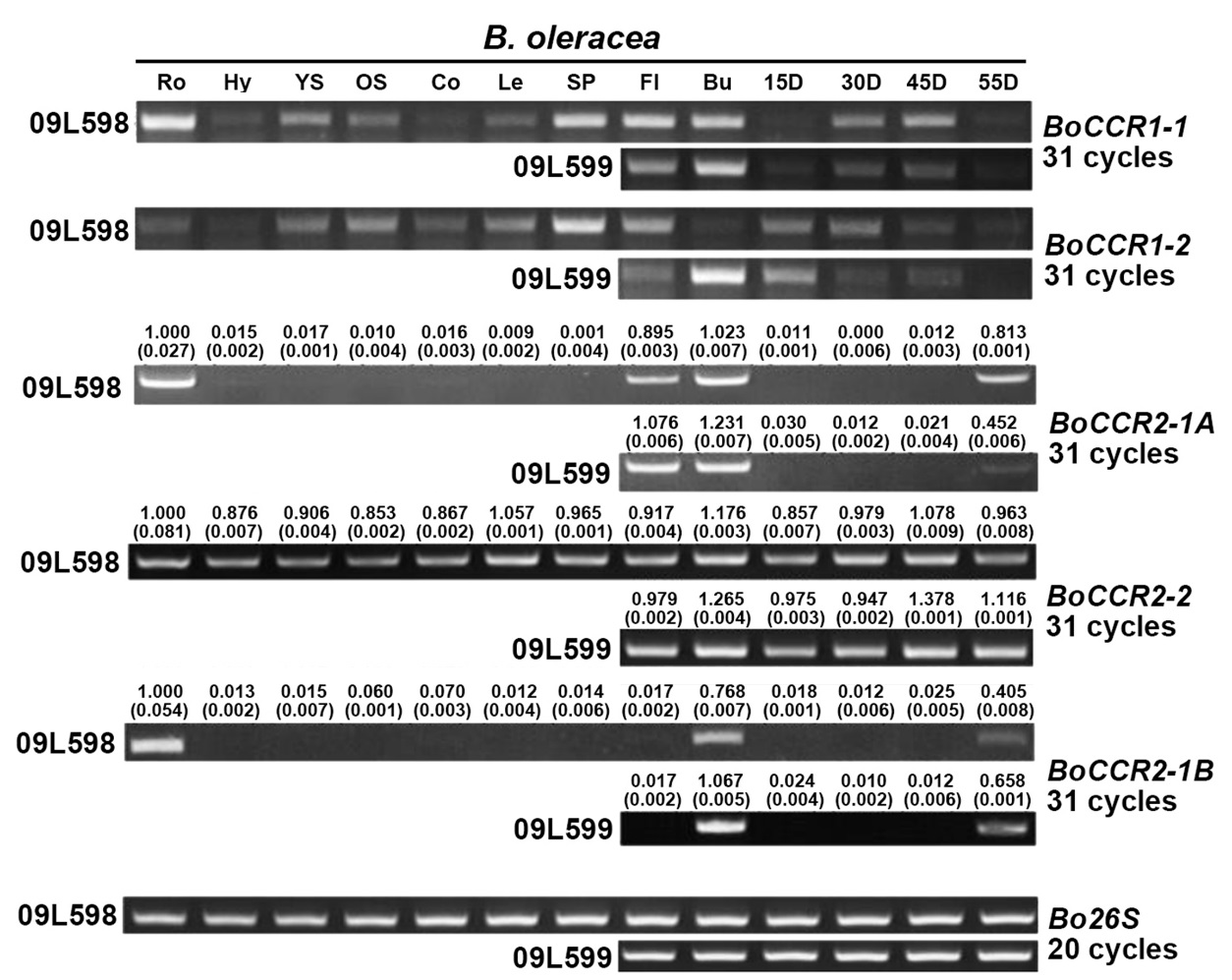

**Fig. S6.** Expression patterns of *CCR1*-genes and *CCR2*-genes show distinct organ-specificity and difference between black and yellow seeds in *B. oleracea.*

09L598 and 09L599 are black- and yellow-seed stocks, respectively. The value (SEM in brackets) above the sRT-PCR band represents the relative expression level of qRT-PCR result. The expression level in root is set as 1.000 for quantification of other organs. Ro, roots; Hy, Hypocotyl; YS, young stems; OS, old stems; Co, cotyledons; Le, leaves; SP, silique pericarp; Fl, flowers; Bu, buds; 15D-55D, seeds of the corresponding days after pollination.

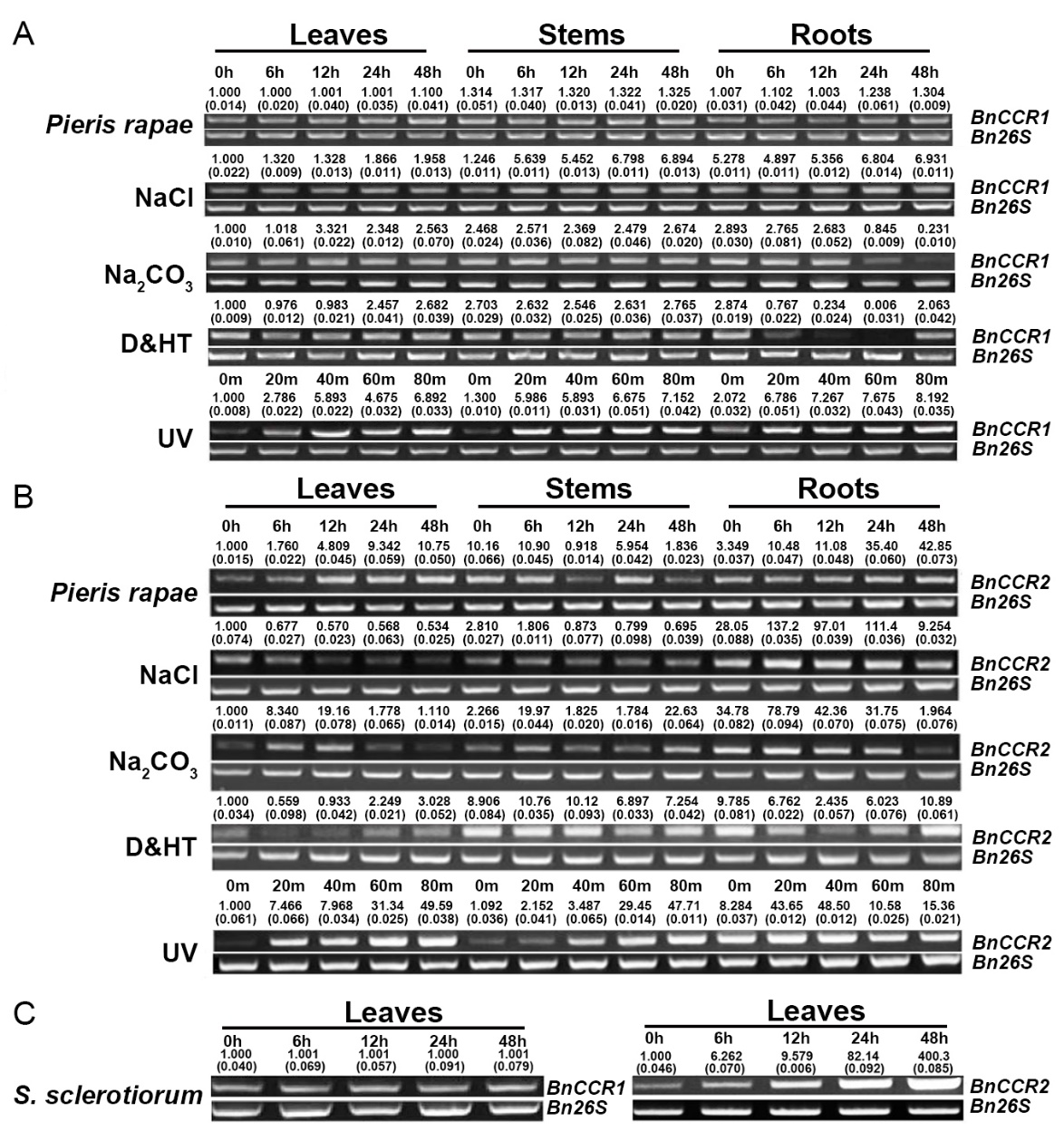

**Fig. S7.** Overall expression of *BnCCR1*-subfamily and *BnCCR2*-subfamily distinctly respond to various stresses in *B. napus* seedlings.

The value (SEM in brackets) above the sRT-PCR band represents the relative expression level of qRT-PCR result. The expression level in the first sample is set as 1.000 for quantification of other samples. m, minutes; D&HT, drought & high temperature.

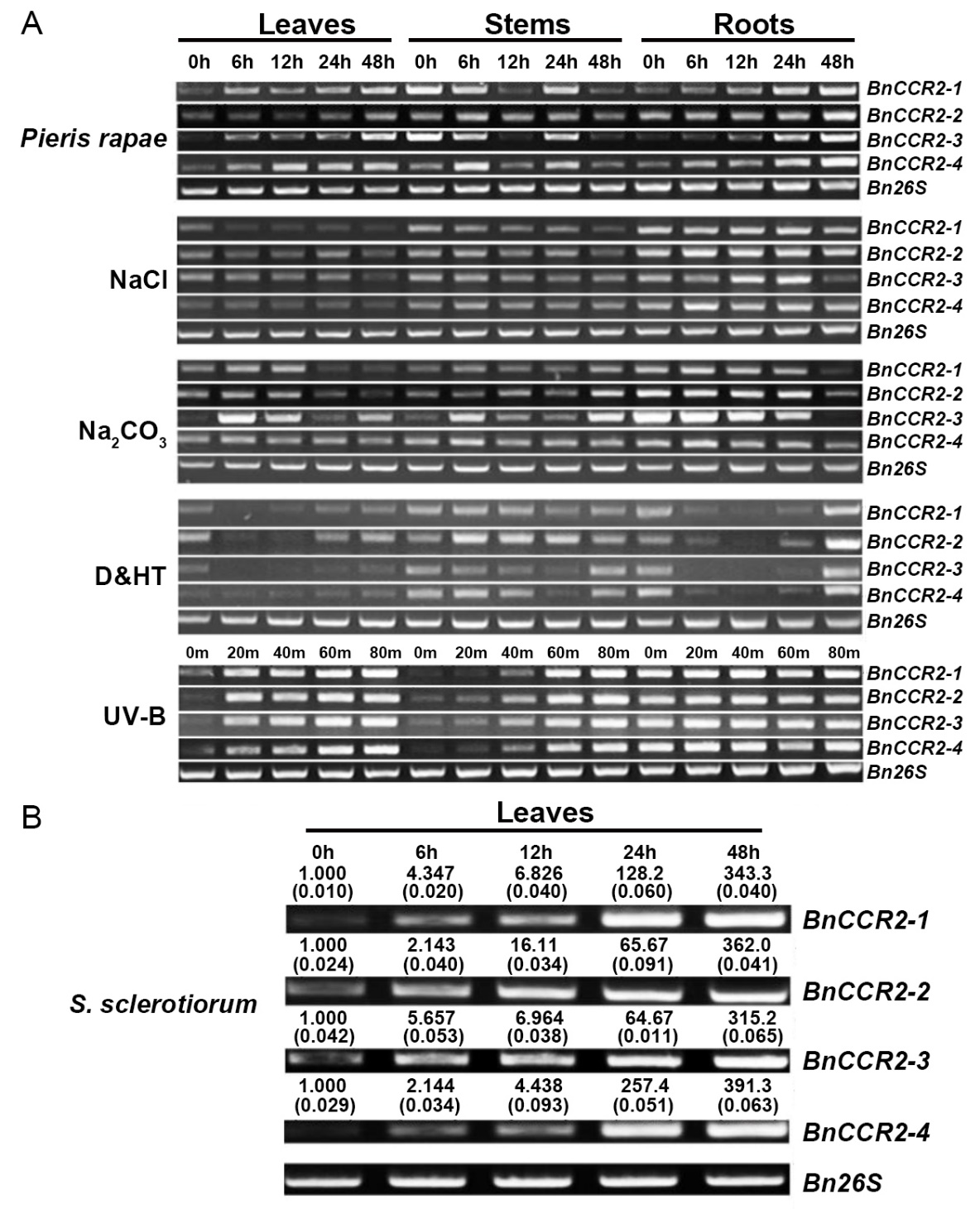

**Fig. S8.** *BnCCR1*-genes and *BnCCR2*-genes distinctly respond to various stresses in *B. napus* seedlings.

The value (SEM in brackets) above the sRT-PCR band represents the relative expression level of qRT-PCR result. The expression level in the first sample is set as 1.000 for quantification of other samples. m, minutes; D&HT, drought & high temperature.

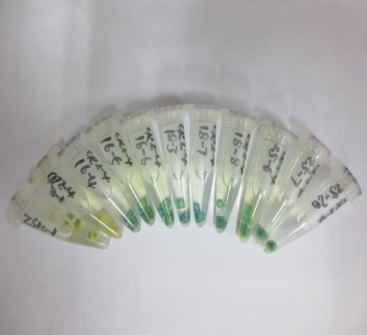

**Basta detection**

**GUS staining**

**PCR detection**

WT

Transgenic lines

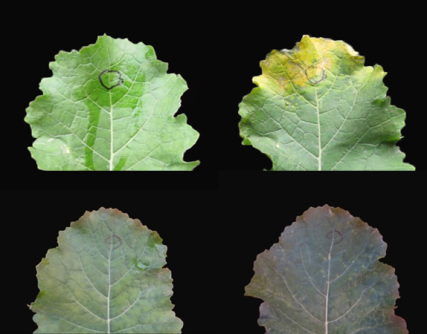

Transgenic lines

WT

Transgenic lines

1^th^ day

7^th^ day

WT

**Fig. S9.** Basta-resistance, GUS-staining and PCR determination of transgenic lines.

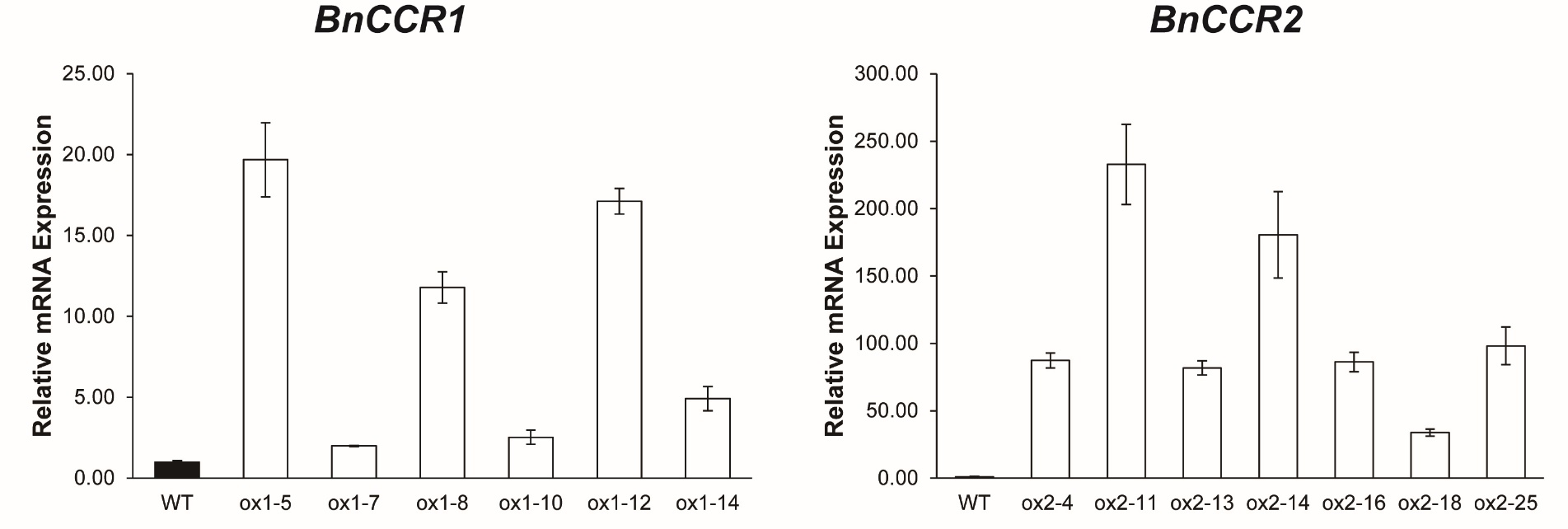

**Fig. S10.** qRT-PCR shows distinct upregulation of the target genes in respective T2 lines of *BnCCR1-2ox* and *BnCCR2-4ox*.

Error bars indicate SD of 3 biological replicates.

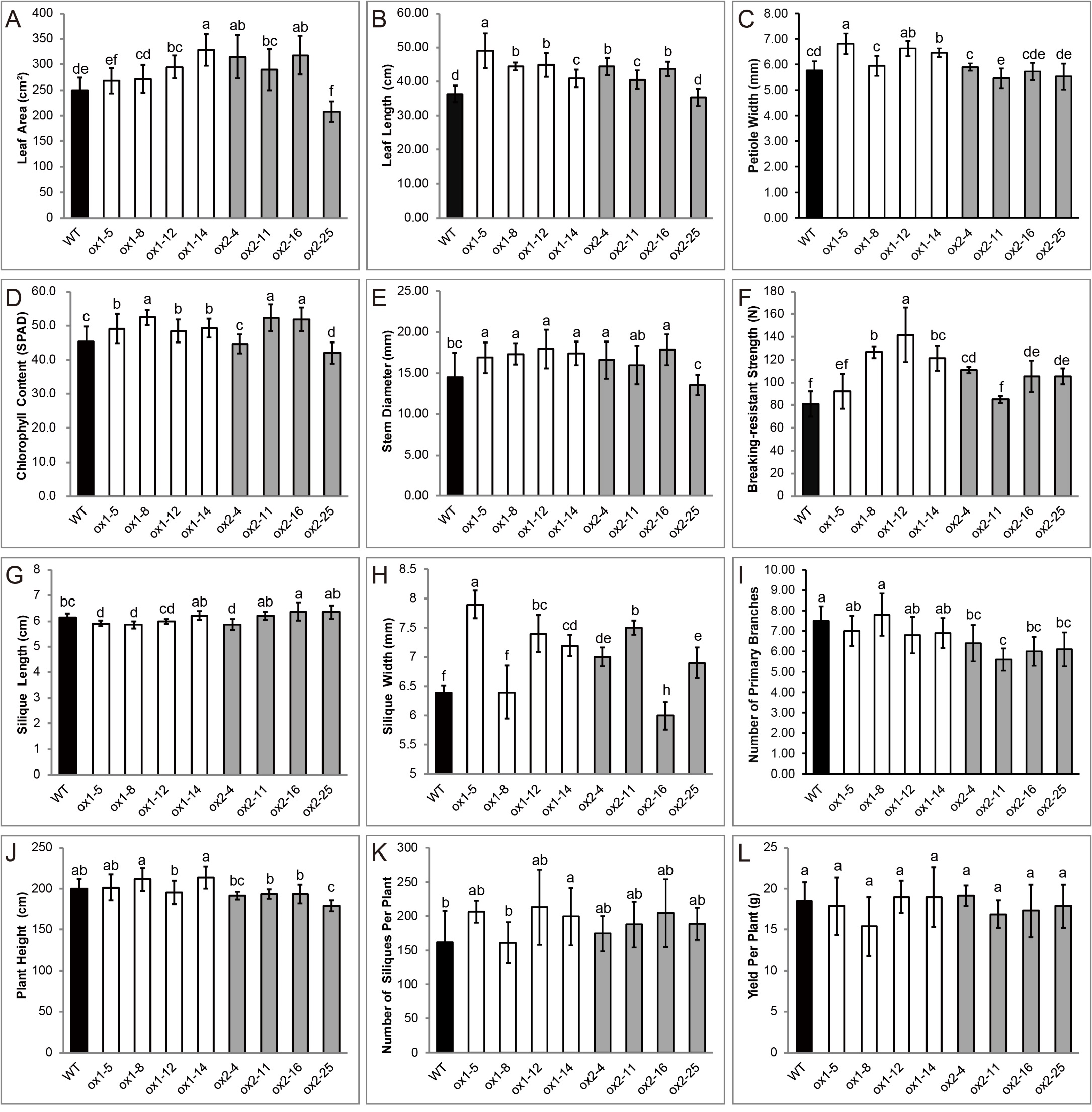

**Fig. S11.** Multiple agronomic traits are significantly modified in *BnCCR1-2ox* and *BnCCR2-4ox* lines.

(A-D) Leaf area (A), leaf length (B), petiole width (C) and chlorophyll content (D) of the fifth and sixth leaves (the most robust leaves) during 9 to 12 leaf-stages.

(E) Diameter of the middle stem at early flowering stage.

(F) Anti-breaking strength of the middle stem at harvest stage after drying.

(G, H) The length and width of the siliques at mature stage.

(I) Number of primary branches.

(J) Plant height at mature stage.

(K) Silique number per plant.

(L) Seed yield per plant.

Data represent means ± SD of at least five biological replicates. Different letters above the bars indicate statistically significant differences (one-way ANOVA, P < 0.05, Duncan’s test).

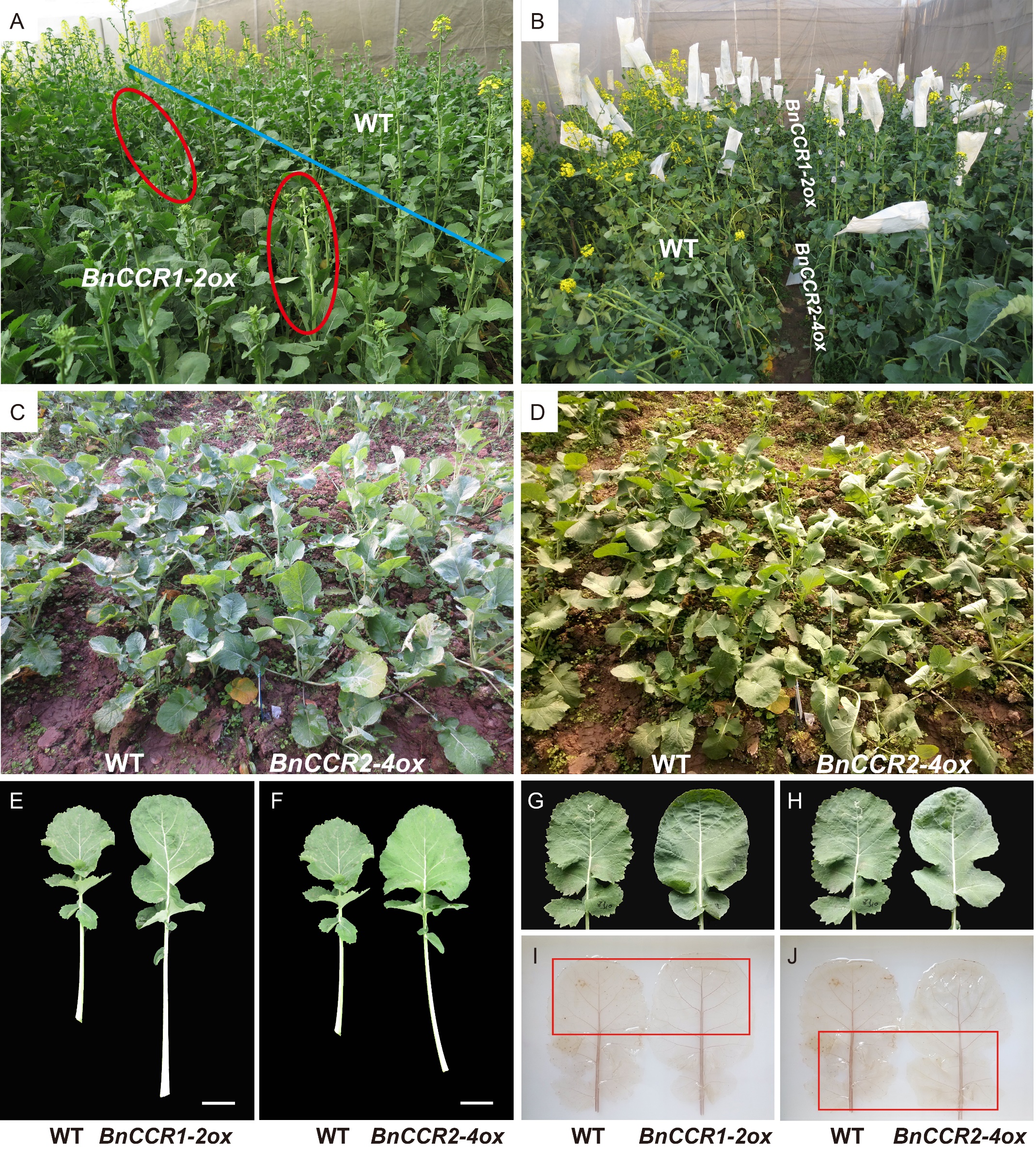

**Fig. S12.** *BnCCR1-2ox* and *BnCCR2-4ox* lines show different growth behaviors and leaf vein strengths.

(A) Delay of bolting and flowering was observed in *BnCCR1-2ox* plants when compared with WT and *BnCCR1-2ox*, and upper stems of *BnCCR1-2ox* at late bolting stage are softer than normal ones and showed a bending phenomenon.

(B) After flowering, both *BnCCR1-2ox* and *BnCCR2-4ox* lines displayed a better lodging resistance than WT, especially for *BnCCR1-2ox* lines.

(C) and (D) Compared with WT and *BnCCR1-2ox*, leaves of *BnCCR2-4ox* seedlings in the morning (about 8:00) have normal angles (C), but showed a rolling-over phenomenon in the afternoon (about 3:00) under strong sunshine (D).

(E, F) At vegetative stage, the strongest leaves of *BnCCR1-2ox* and *BnCCR2-4ox* were distinctly larger than WT, both had similar effect on leaf blade growth, but *BnCCR1-2ox* had larger effect on petiole-vein growth than *BnCCR2-4ox*.

(G-J) Though visual observation of petiole-vein system of both *BnCCR1-2ox* and *BnCCR2-4ox* were larger than WT ([G] and [H]), phloroglucinol-HCl staining showed that petiole-vein lignification was strengthened only in *BnCCR1-2ox* plants while weakened in *BnCCR2-4ox* plants ([I] and [J]).

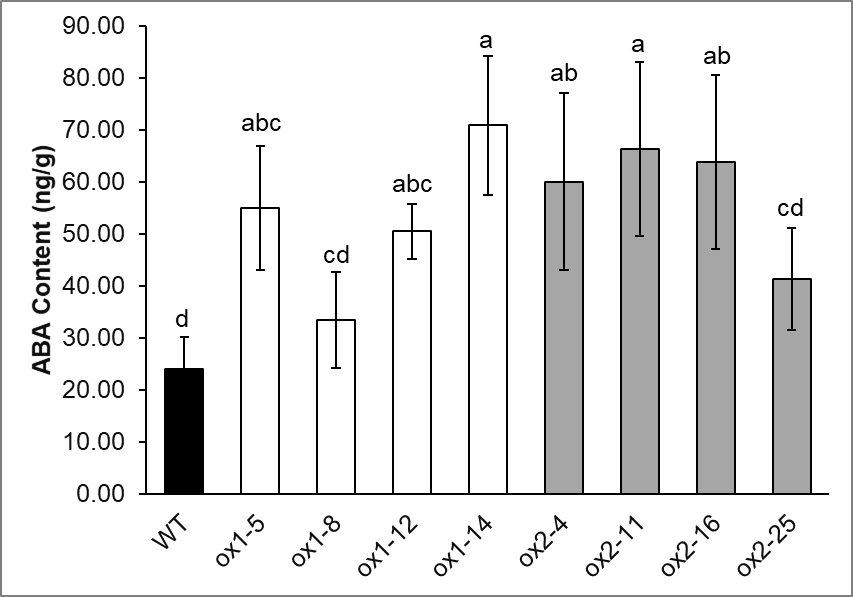

**Fig. S13.** ABA content is upregulated in leaves of *BnCCR1-2ox* and *BnCCR2-4ox* lines.

Data represent means ± SD of at least 3 biological replicates. Different letters above the bars indicate statistically significant differences (one-way ANOVA, P < 0.05, Duncan’s test).

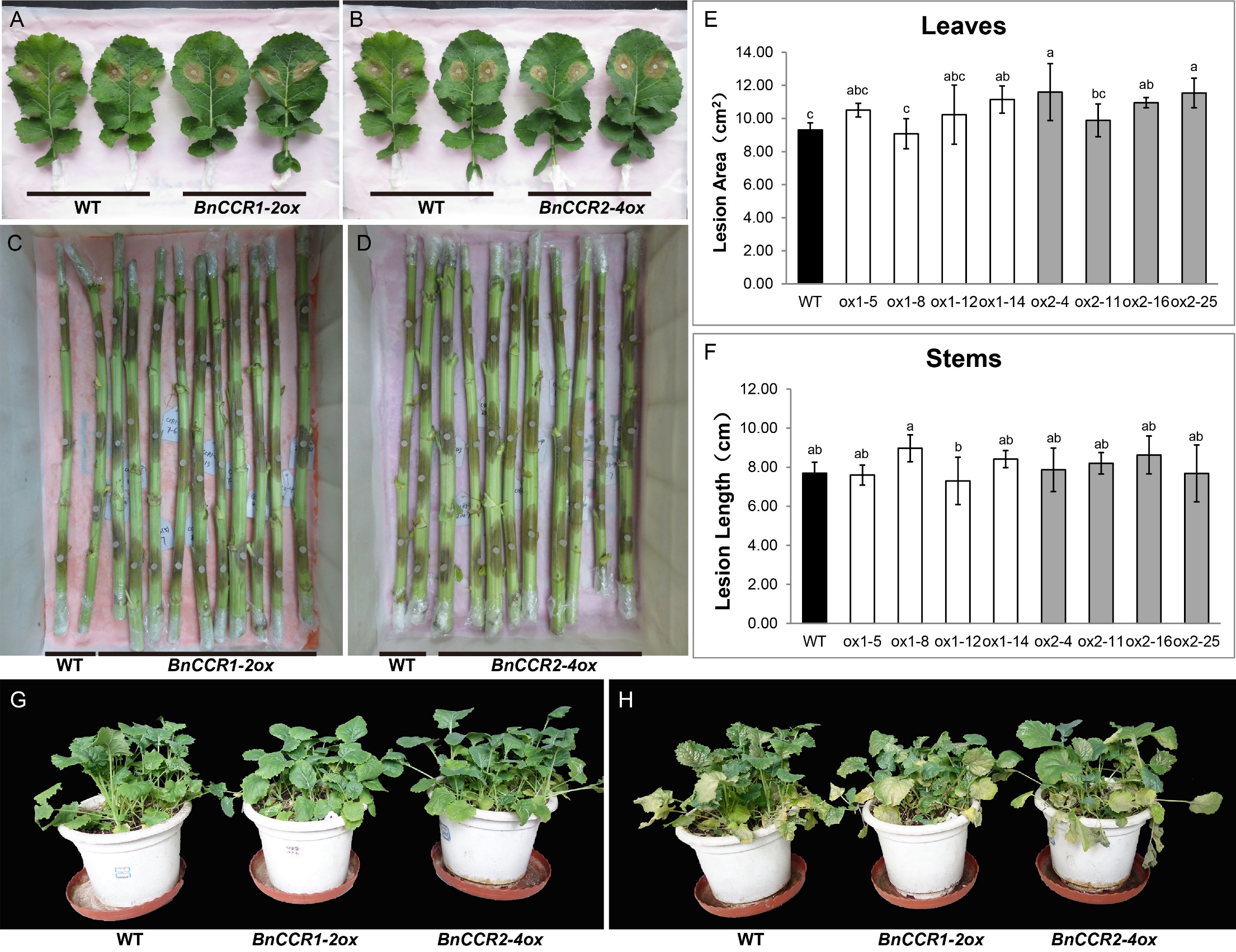

**Fig. S14.** Plants of both *BnCCR1-2ox* and *BnCCR2-4ox* do not show enhanced resistance to *S. sclerotiorum* and UV-B.

(A-D) Lesions on detached leaves and stems after 48 h and 72 h of inoculation respectively.

(E, F) Lesion areas on leaves and lesion lengths on stems respectively. Data represent means ± SD of at least 3 biological replicates. Different letters above the bars indicate statistically significant differences (one-way ANOVA, P < 0.05, Duncan’s test).

(G, H) Pot seedlings before UV-B treatment and 1 week after 1-h UV-B treatment respectively.

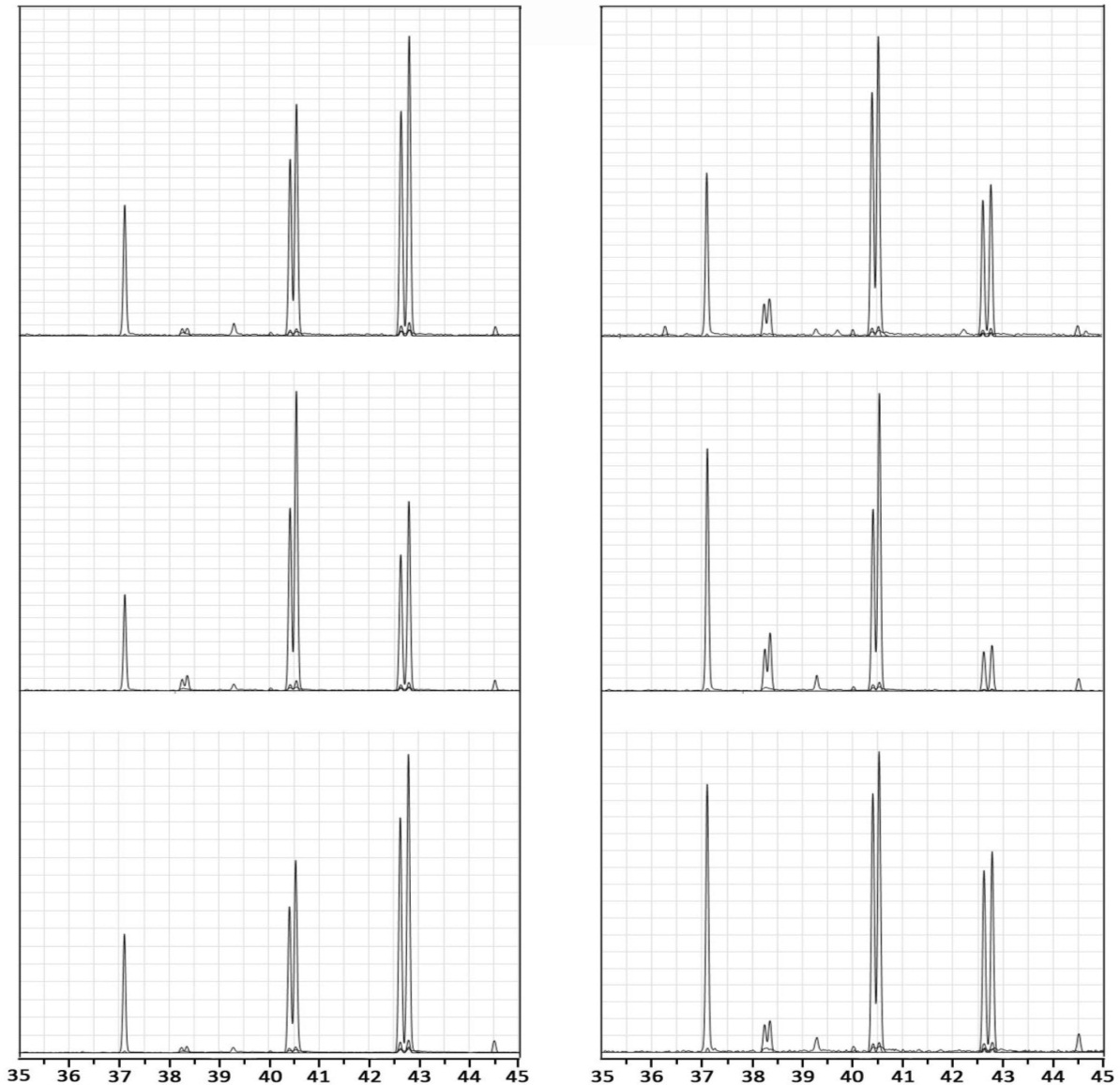

**IS**

**H1**

**G1**

**G2**

**S1**

**S2**

Relative abundance

Retention time（min）

WT stem

WT root

ox1-5 stem

ox1-5 root

ox2-16 stem

ox2-16 root

**H2**

**Fig. S15.** GC-MS chromatograms show different changes of monolignol proportions in *BnCCR1-2ox* and *BnCCR2-4ox* lines.

IS, Internal standard tetracosane; H1, H2, G1, G2, S1 and S2 represent the isomers of the corresponding H-, G- and S-lignin monomers.

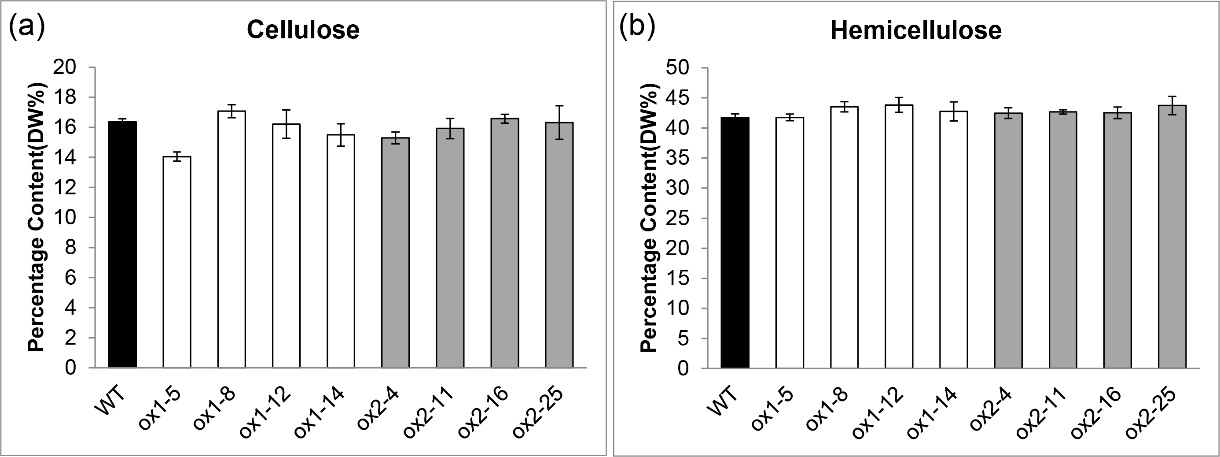

**Fig. S16.** Contents of cellulose and hemicellulose in the stems of *BnCCR1-2ox* and *BnCCR2-4ox* lines are not changed based on NIRS detection.

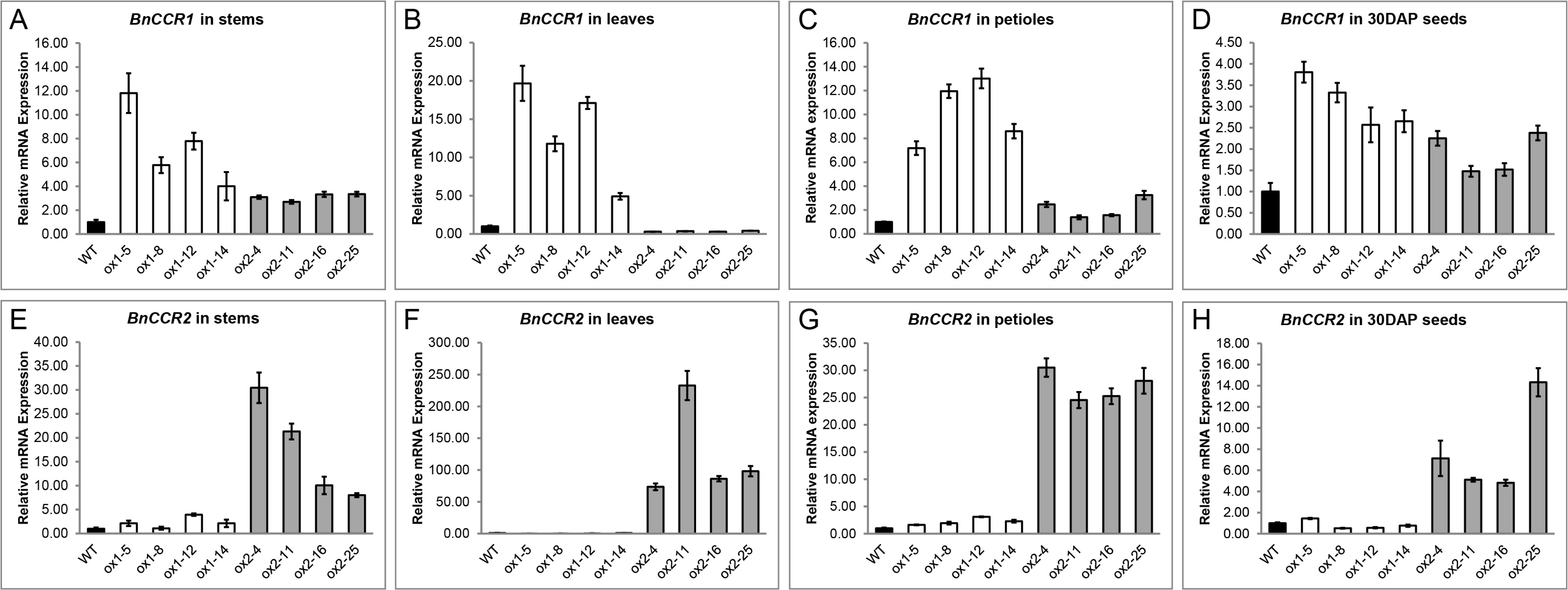

**Fig. S17.** *BnCCR1* expression is obviously altered in *BnCCR2-4ox* lines, whereas *BnCCR2* expression has little change in *BnCCR1-2ox* lines.

(A-D) *BnCCR1* expression is upregulated in stems, petioles and seeds, while downregulated in leaves, of *BnCCR2-4ox* lines.

(E-H) *BnCCR2* expression is almost not changed in stems, leaves, petioles and seeds of *BnCCR1-2ox* lines.

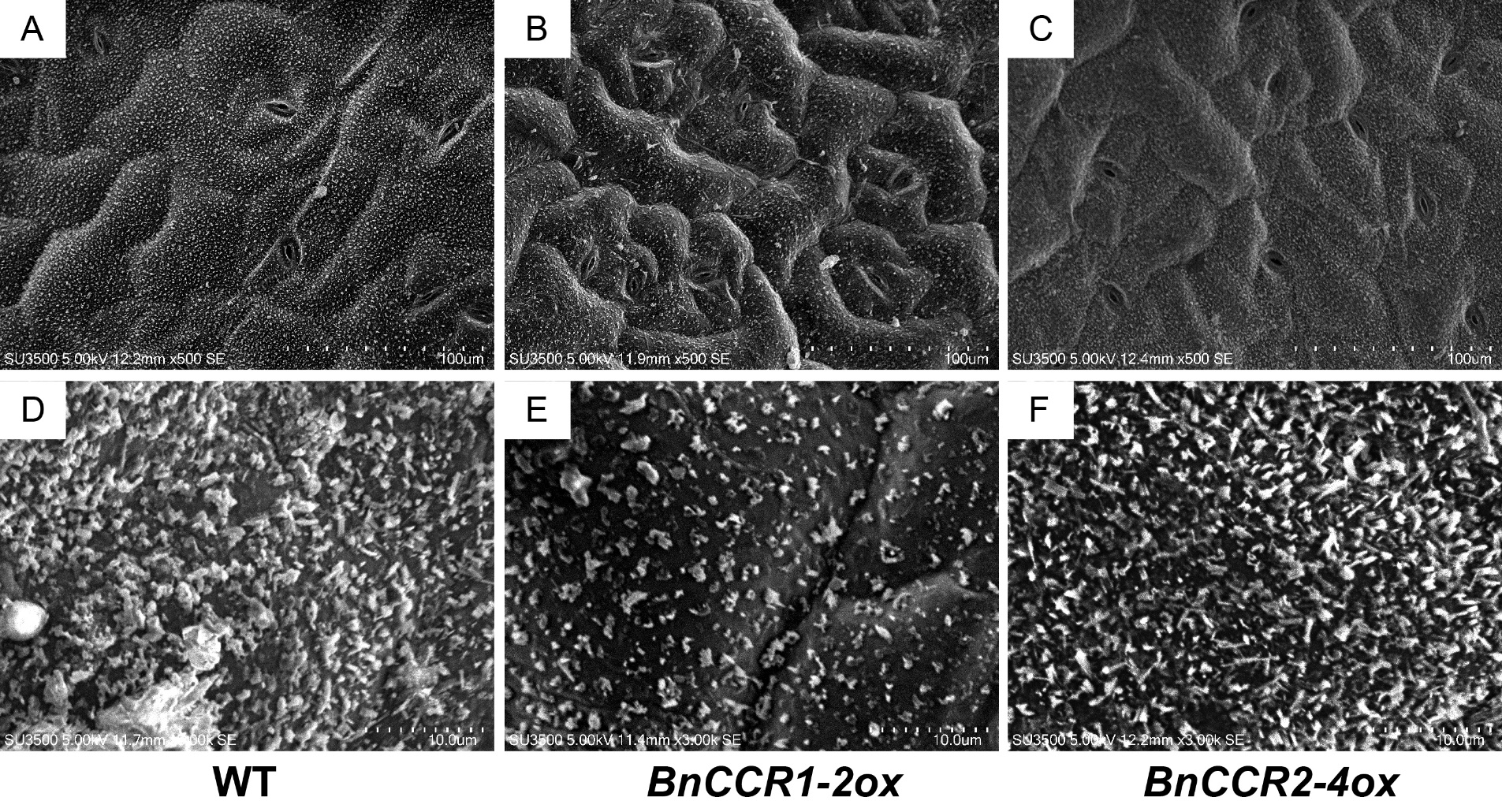

**Fig. S18.** Leaf surfaces of *BnCCR1-2ox* and *BnCCR2-4ox* plants are less flat with decreased wax deposition than WT.

(A) to (C): Electron microscopy observation shows that leaf surface cells of *BnCCR1-2ox* and *BnCCR2-4ox* plants are more prominent than WT. Bar = 100 μm.

(D) to (F): Electron microscopy observation shows that leaf epidermal wax crystals of *BnCCR1-2ox* and *BnCCR2-4ox* plants are less than WT. Bar = 10 μm.

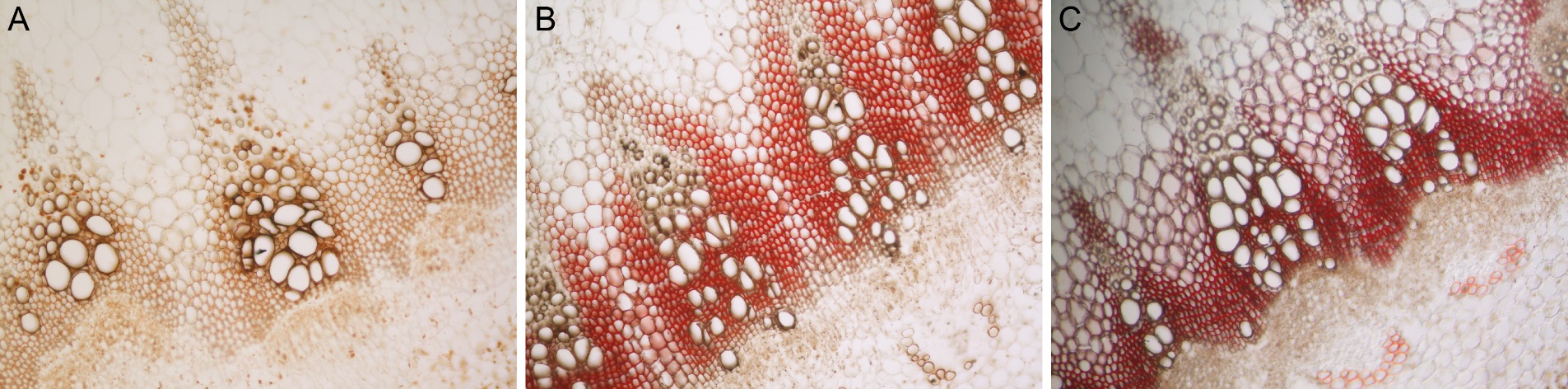

**Fig. S19.** Mäule staining of stem sections indicates a trend of increase of S-type lignins during developmental process in *B. napus*.

(A) Middle-upper part of the stem at early flowering stage.

(B) Middle-lower part of the stem at early flowering stage.

(C) Middle-upper part of the stem at mature stage.
