## Supplementary material for "Two Types of Cinnamoyl-CoA Reductase Function Divergently in Tissue Lignification, Phenylpropanoids Flux Control, and Inter-pathway Cross-talk with Glucosinolates as Revealed in *Brassica napus*": Supplymentary Methods-20210413

**Supplementary Methods**

**Stress** **treatments**

***S. sclerotiorum* infection and resistance investigation.** The *S. sclerotiorum* strain was kindly provided by Prof. Wei Qian (Southwest University, China), and the inoculation on *B. napus* was performed as described previously (Mei *et al.*, 2011). The 4th or 5th fully expanded leaves at the 9- to 12-leaf stage and stem segments at early flowering stage were sampled for resistance assay in the laboratory. Moist towels were placed at the bottom of an appropriate container. Filter papers were placed over the towels, and the detached leaves and stem segments were placed on the filter papers. Two plugs (6 mm in diameter) punched from the growing margin of 3-day-old culture of *S. sclerotiorum* on PDA medium were placed separately on two sides of the midrib of leaf and the stem segment. The container was sealed with plastic film to keep humidity, with temperature maintained at 21°C in laboratory. The lesion size (S) was calculated with the formula S=π×a×b/4, where a and b represent the long and the short diameters of the ellipse-like lesion which was measured 2 days after leaf inoculation or 3 days after stem inoculation. Leaf discs of 2-3 cm in diameter around the infection site were sampled after 0, 6, 12, 24 and 48 h of inoculation.

***P. rapae*** **inoculation treatment.** Leaves of the same position of normal-growth *B. napus* seedlings in pots were inoculated with *P. rapae* worms of similar size, which were laid on *B. napus* plants in preparation. Each leaf was inoculated with 4 worms, and each inoculated plant was covered with a plastic net to prevent worm escaping. Leaves, stems and roots were sampled after 0, 6, 12, 24 and 48 h of inoculation.

**Treatment with high temperature & drought.** Normal-growth *B. napus* seedlings in pots were put into a growth chamber which was kept at 37°C with no watering. Leaves, stems and roots were sampled after 0, 6, 12, 24 and 48 h of treatment.

**Salt treatment.** Normal-growth *B. napus* seedlings in pots were watered with 500 mL NaCl or Na_2_CO_3_ (0.3 mol/L), and the leaves, stems and roots were collected after 0, 6, 12, 24 and 48 treatment.

**UV-B Treatment.** Normal-growth *B. napus* seedlings in pots were irradiated with UV-B light from 4 UV-B lamps (2.4 mW/cm^2^), and leaves, stems and roots were collected after 0, 20, 40, 60 and 80 min of treatment.

**Isolation of *Brassica* *CCR* cDNA and Genomic sequences**

All samples were frozen in liquid nitrogen and stored at -80°C. Total RNA was extracted from roots, stems, leaves, flowers and developing seeds of the 3 *Brassica* species using EASYspin Kit (Biomed, China) and RNAprep pure plant kit (TIANGEN, China). Equal 1 µg RNA from each organ were pooled for total cDNA synthesis for 5’-RACE and 3’-RACE using GeneRacer^TM^ Kit (Invitrogen, USA). Based on *in-silico* cloning and multi-alignment of *Brassica* genomic and cDNA tags from NCBI GenBank nr/nd, GSS and EST databases, RACE (rapid amplification of cDNA ends) primers were designed according to conservative sites of *CCR* genes (Supplementary Table S8).

5’-RACE primary and nested amplifications of *Brassica CCR1* genes were performed using primer pairs 5’P+RCCR1 and 5’NP+RCCR1-1N / 5’NP+RCCR1-2N, while 3’-RACE primary and nested amplifications of *Brassica CCR1* genes were performed using primer pairs 3’P+FCCR1C1 and 3’NP+FCCR1C2, respectively. 5’-RACE primary and nested amplifications of *Brassica CCR2* genes were performed using primer pairs 5’P+RCCR2-51 and 5’NP+RCCR2-52, while 3’-RACE primary and nested amplifications of *Brassica CCR2* genes were performed using primer pairs 3’P+FCCR2-31 and 3’NP+FCCR2-32, respectively. All reactions were performed using standard Taq-PCR, annealed at 52°C, with extension time of 1 min.

The full-length cDNAs of *Brassica CCR* genes were PCR-amplified from the 3’-RACE total cDNA template using Taq plus enzyme (a mixture of Pfu proofreading and Taq DNA polymerases). The primers are listed in Supplementary Table S8. Forward primers FCCR1-1, FCCR1-2, FBCR1-13 and FBCR1-24 were paired with reverse primers RCCR1-1, RCCR1-2, RBCR1-3, RBCR1-1 and RBCR1-24 to form 20 primer combinations to amplify full-length cDNAs of *CCR1* subfamily genes from *B. napus*, *B. rapa* and *B. oleracea*. For amplification of *CCR2* subfamily genes from the 3 *Brassica* species, forward primers FBNCR210, FBRCR221, FBOCR2E1 and FBOCR2G5 were paired with reverse primers RBNCR22, RBNCR28, RBOCR22 and RBOCR2G9 to form 16 primer combinations. Gradient annealing temperature of 52-60°C was adopted for each PCR, with an extension time of 2 min. All successful primer pairs were used to amplify the corresponding gDNA sequences by replacing the total cDNA template with total gDNA of respective species. *Brassica CCR2* pseudogenes were directly amplified without RACE information using corresponding primers listed in Supplementary Table S8. PCRs for gDNA and pseudogenes had an annealing temperature of 55°C and an extension time of 3 min.

Agarose gel electrophoresis, gel recovery, TA-cloning with pMD 19-T vector (TaKaRa Dalian, China), *Escherichia coli* transformation, colony identification and Sanger’s sequencing were performed with regular molecular methods.

**Bioinformatic Methods for this Study**

Open reading frame (ORF) finding and translation, parameter calculation and sequence alignment of genes and proteins were performed on Vector NTI Advance 11.53 (Invitrogen, USA). Structural and functional predictions of proteins were performed on Expasy website (http://www.expasy.org).

After our cloning work of *Brassica* *CCR* family, whole genome sequencing and annotation data of *B. rapa*, *B. oleracea* and *B. napus* were available in GenBank, then we performed *in-silico* cloning of all *CCR1* and *CCR2* subfamily genes from these 3 species. Using the cloned *Brassica* *CCR* sequences as queries (E-probes), BLASTn and BLASTp were performed against the nr/nt, nr refseq_rna, refseq_protein, refseq_genomes, wgs, est and tsa databases of *Brassica* (taxid:3705) on NCBI (https://blast.ncbi.nlm.nih.gov/Blast.cgi), Genoscope (http://www.genoscope.cns.fr/brassicanapus/) and BRAD (http://39.100.233.196/#/), sequences of all *CCR1* and *CCR2* genes and proteins from *Brassica* were *in-silico* cloned and multi-aligned with the actually cloned sequences in this study.

For phylogenetic analysis, all true-CCR protein sequences from representative whole-genome-sequenced species of Brassicales and other malvids orders were *in-silico* cloned into a fasta file, which was sent to Seaview 4.0 to perform clustalo multi-alignment and construct a Squared-type phylogenetic tree using the Distance-BioNJ-Poisson mode. The reliability of the tree was measured by bootstrap analysis with 1,000 replicates.

**Construction of Overexpression Vectors**

The *BnCCR1-2* and *BnCCR2-4* full-length cDNAs harbored on the sequenced recombinant pMD 19-T vectors were subcloned into platform vector pC2301M1DPB using *Bam*HI+*Sac*I double digestion and *Xba*I single digestion, under the control of a double CaMV35S promoter (P_D35S_) and the nopaline synthase terminator (T-nos) (Fu *et al.*, 2017), to generate plant overexpression vectors pCD-*BnCCR1-2ox* and pCD-*BnCCR2-4ox*, respectively. The overexpression vectors were transformed into *Agrobacterium tumefaciens* (LBA4404) through a freeze–thaw method (Chen *et al.*, 1994), and engineering strains were obtained after Taq-PCR verification (annealed at 55°C, extension for 1.5 min) with primer pairs F35S3N+RCCR1-2 and FCCR1-2+RNOS5N for pCD-*BnCCR1-2ox* and primer pairs F35S3N+RBNCR22 and FBNCR210+RNOS5N for pCD-*BnCCR2-4ox*.

**Measurement of Leaf SPAD Readings**

A SPAD-502 chlorophyll meter (Minolta, Osaka, Japan) was used to obtain SPAD values (SPAD units) from the 4th or the 5th fully expanded leaves of five representative transgenic plants of each line. Three average distributed points SPAD readings were obtained per leaf, and averaged as the mean SPAD reading of the leaves (Yang *et al.*, 2014).

**Near-infrared Reflectance Spectroscopy (NIRS) Measurements**

Measurements for cellulose and hemicellulose of the stem, seed color, oil content, fatty acid profile, total glucosinolates and fibre components of the seeds were obtained using an NIR System 6500 with WinISI II software (FOSS GmbH, Rellingen, Germany) according to (Liu *et al.*, 2012). The dried harvested stems were ground with a universal mill to pass a 40-mesh screen. The calibrations which were developed by our laboratory specifically for the measurement of these traits for *B. napus* seeds and stems, were used to detect the phenotype values for seed colour (visual light absorbance), acid detergent lignin (ADL, % dry weight), acid detergent fibre content (ADF, % dry weight) and neutral detergent fibre content (NDF, % dry weight) from NIRS spectra. NIRS-derived estimates for each trait and genotype were averaged over two technical repetitions. Cellulose concentrations were calculated as the difference between NDF (predominantly phenylpropanoid compounds, cellulose and hemicellulose) and ADF (predominantly phenylpropanoid compounds and hemicellulose), and hemicellulose concentration as the difference between ADF and ADL (phenylpropanoids).

**Chen H, Nelson RS, Sherwood JL**. 1994. Enhanced recovery of transformants of Agrobacterium tumefaciens after freeze-thaw transformation and drug selection. Biotechniques **16**, 664-668, 670.

**Fu C, Chai YR, Ma LJ, Wang R, Hu K, Wu JY, Li JN, Liu X, Lu JX**. 2017. Evening primrose (*Oenothera biennis*) Δ6 fatty acid desaturase gene family: cloning, characterization, and engineered GLA and SDA production in a staple oil crop. Molecular Breeding **37**(6), 83.

**Liu LZ, Stein A, Wittkop B, Sarvari P, Li JN, Yan XY, Dreyer F, Frauen M, Friedt W, Snowdon RJ**. 2012. A knockout mutation in the lignin biosynthesis gene *CCR1* explains a major QTL for acid detergent lignin content in *Brassica napus* seeds. Theoretical and Applied Genetics **124**, 1573-1586.

**Mei J, Qian L, Disi JO, Yang X, Li Q, Li J, Frauen M, Cai D, Qian W**. 2011. Identification of resistant sources against *Sclerotinia sclerotiorum* in *Brassica* species with emphasis on *B. oleracea*. Euphytica **177**, 393-399.

**Yang H, Li JW, Yang JP, Wang H, Zou JL, He JJ**. 2014. Effects of nitrogen application rate and leaf age on the distribution pattern of leaf spad readings in the rice canopy. Plos One **9**(6), 83.
